# Multi-layered proteomics identifies insulin-induced upregulation of the EphA2 receptor via the ERK pathway and dependent on low IGF1R level

**DOI:** 10.1101/2023.12.14.571674

**Authors:** Sarah Hyllekvist Jørgensen, Kristina Bennet Emdal, Anna-Kathrine Pedersen, Lene Nygaard Axelsen, Helene Faustrup Kildegaard, Damien Damozay, Thomas Åskov Pedersen, Mads Grønborg, Rita Slaaby, Peter Kresten Nielsen, Jesper Velgaard Olsen

## Abstract

Insulin resistance impairs the cellular insulin response and frequently precedes metabolic disorders, like type 2 diabetes, which are affecting an increasing number of people globally. Given the critical role of the liver in glucose and lipid metabolism, understanding the molecular mechanisms in hepatic insulin resistance is essential for early preventive treatments. To elucidate changes in insulin signal transduction associated with hepatocellular resistance, we employed a multi-layered mass spectrometry-based proteomics approach focusing on insulin receptor (IR) signaling at the interactome, phosphoproteome, and proteome levels.in a long-term hyperinsulinemia-induced insulin-resistant HepG2 cell line with a knockout of the insulin-like growth factor 1 receptor (IGF1R KO). Analysis of the dynamic insulin-induced IR interactome revealed recruitment of the PI3K complex in both insulin-sensitive and -resistant cells. From the phosphoproteomics dataset, a change in insulin-stimulated signaling responses in insulin resistance was observed and showed attenuated signaling via the metabolic PI3K-AKT pathway but sustained extracellular signal-regulated kinase (ERK) activity. At the proteome level, the ephrin type-A receptor 2 (EphA2) showed an insulin-induced increase in expression. This receptor belongs to the Eph receptor family and participates in various cellular processes, such as cell adhesion, migration, and tissue development. The protein abundance regulation of EphA2 occurred through the ERK signaling pathway and was concordantly independent of insulin resistance. Induction of EphA2 by insulin was confirmed in other cell lines and observed uniquely in cells with high levels of IR compared to IGF1R. The multi-layered proteomics dataset provided insights into insulin signaling in general and in the context of insulin resistance, and it can going forward serve as a resource to generate and test hypotheses, leading to an improved understanding of insulin resistance.

## Introduction

Insulin resistance, a hallmark of type 2 diabetes, is characterized by impaired cellular responses to insulin [1], [2]. It affects nearly 40% of young American adults [3] and is influenced by various factors, including obesity, physical inactivity, chronic inflammation, prolonged exposure to elevated levels of nutrients, and pathophysiological insulin levels [4], [5]. The cellular response and mechanisms of insulin resistance vary by tissue, with fat, muscle, and liver being the primary insulin-responsive tissues [1].

The liver plays a key role in glucose homeostasis and overall metabolism, and insulin signaling in hepatocytes primarily coordinates glucose uptake, storage, and production [6]. Insulin binds to the insulin receptor (IR), a plasma membrane-spanning receptor tyrosine kinase (RTK), leading to its autophosphorylation. Insulin signaling branches into two major pathways - a metabolic pathway via the phosphatidylinositol 3-kinase (PI3K)-AKT signaling axis, which plays the primary role in hepatic insulin response, and a mitogenic pathway controlling cell growth, proliferation, and gene expression [7]–[10]. The growth factor capabilities of insulin depend on signaling activation via the canonical extracellular signal-regulated kinase (ERK) pathway [8]. These central signaling pathways have several shared and diverging points of regulation by various kinases and phosphatases, leading to complex interplay and cross-talk in downstream activation [11], [12]. Hepatic insulin resistance primarily impairs PI3K-AKT signaling [2]. The relationship between IR levels and insulin resistance is unclear, with some studies reporting reduced IR levels in insulin-resistant states [13]–[16] and others observing no significant differences between insulin-sensitive and insulin-resistant groups [17], [18]. Studies have shown that changes in subcellular localization of IR may play a role in hepatic insulin resistance [17].

Quantitative mass spectrometry (MS)-based proteomics is a powerful technology for global analysis of cell signaling networks, and it is widely used to characterize signaling pathways from activated RTKs in different cell models [19]–[22]. MS-based interactome studies have used receptor overexpressing cell models to study the proximal signaling of the IR [22]–[24]. While occasionally applied in phosphoproteomics, overexpression can lead to aberrations in interactions, downstream signaling, and general cellular processes. Additionally, induction of insulin resistance in such manipulated cell line models is often more complicated or unattainable [22]. To gain a deeper understanding of the molecular mechanisms underlying insulin resistance, we performed a multi-layered MS-based proteomics investigation in a HepG2 IGF1R KO cell line model with endogenous IR expression. The study focused on analyzing the insulin-dependent IR interactome, phosphoproteome, and proteome to uncover molecular changes driving insulin resistance in the model. The global phosphoproteome analysis affirmed that the insulin-resistant condition pathway specifically reduced PI3K-AKT response but not the ERK pathway. The proteome analysis revealed upregulated protein levels of the RTK, ephrin type-A receptor 2 (EphA2), in insulin-resistant compared to -sensitive cells. EphA2 belongs to the Eph receptor family and is involved in diverse cellular processes, including cell adhesion, migration, and tissue development [25], [26]. Moreover, the receptor displays upregulation in certain types of cancer [26]–[28]. This study showed that the activation of the ERK pathway by insulin was responsible for upregulating EphA2 levels, and this mechanism was independent of insulin resistance. The ERK pathway dependency was shown by MEK inhibition and the lack of induction with the AKT-biased partial IR agonist, S597 [29], [30]. Screening of different cell lines suggested that the ability of insulin to induce EphA2 protein levels required a high IR/IGF1R ratio. Exploring this previously unreported regulation of EphA2 by insulin and its correlation with the relatively higher IR/IGF1R ratio has the potential to provide insights into the intricate insulin signaling in diverse cellular and tissue contexts.

## Methods

### Experimental Design and Statistical Rationale

To investigate molecular changes associated with insulin resistance, we utilized an insulin-like growth factor 1 receptor knock-out (IGF1R KO) HepG2 hepatocellular carcinoma cell line. This cell model offers an advantage over the commonly used wild-type (WT) HepG2 cell line, which expresses IGF1R at comparable levels to IR, not typically observed in healthy hepatocytes [31], [32]. Employing the IGF1R KO cell line allowed a focused characterization of the IR-mediated insulin response, disregarding the influence of insulin signaling through IR-IGF1R hybrids. An insulin-resistant state was induced by hyperinsulinemia based on a protocol established by Dall’Agnese *et al*. [17], in which the cells were stimulated with 0.1 nM (considered physiological) or 3 nM (considered pathological) insulin for 48 hours. To ensure comprehensive coverage of differential insulin signaling pathways, the proteome, phosphoproteome, and IR interactome of insulin-sensitive and -resistant cells were sampled unstimulated or after 5-minute insulin stimulation with 0.1, 3, and 100 nM insulin. Five biological replicates were collected for each experimental condition. The same co-immunoprecipitation (co-IP) buffer-extracted lysates were used to generate all three types of datasets. Also, a single-shot proteome dataset was acquired using cells lysed in SDS lysis buffer. Validation of the insulin-resistant phenotype involved the application of several orthogonal methods. This comprehensive experimental design and analysis strategy enabled a thorough exploration of insulin signaling dynamics in an insulin-sensitive and -resistant state. Data from a stable isotope labeling of amino acids in cell culture (SILAC) experiment in the rat hepatoma cell line, H4IIE, was included, analyzing proteome changes after 24- and 48-hour treatment with insulin or the IR agonist S597 [29], [30], using three biological replicates. Additionally, a SILAC H4IIE phosphoproteome dataset after stimulation with insulin or S597 for 5, 15, and 30 minutes was acquired, including two biological replicates. Details on the bioinformatics and statistical approach for different data analyses are provided in the relevant sections. The number of replicates used for the analyses, aside from the MS data, is specified in the respective figure legends and sections.

### Reagents

Human insulin (Novo Nordisk A/S) and S597 [30] (Novo Nordisk A/S) was utilized for the experiments. The following antibodies were applied for Western blots (WBs): rabbit anti-phospho-EPHA2 (Tyr594 and S897), rabbit anti–phospho–IGF-1Rβ (Tyr1131)/insulin receptor β (Tyr1146), rabbit anti–insulin receptor β (4B8), rabbit anti–phospho-Akt (Thr308) and anti-Akt, mouse anti– phospho-ERK1/2 (Thr202/Tyr204), rabbit anti-ERK1/2 (Cell Signaling Technology); rabbit anti-Eph receptor A2 antibody (ab273118 and ab185156), mouse anti-beta actin antibody (ab8226) (Abcam); insulin receptor β antibody (CT-3) (Santa Cruz); goat anti-rabbit immunoglobulin (IgG) secondary antibody, horseradish peroxidase (HRP)–conjugated, and goat anti-mouse IgG secondary antibody, HRP-conjugated (Bio-Rad). For phospho (p)Tyr enrichment in the SILAC experiment, pTyr1000 (8803) and pTyr100 (5636) (Cell Signaling Technology) were used. For immunofluorescence and co-IP, the mouse-anti insulin receptor (83-7) obtained under license from Professor K. Siddle, University of Cambridge, UK [33] was used. The secondary antibody was Alexa Fluor 488 goat anti-mouse IgG (A-11001) (Thermo Fisher Scientific). For flow cytometry analysis, the following antibodies were used: anti IR (D2) mouse IgG110 [34] and mouse IgG1 Neg control (DAKO). The kinase inhibitors used were cobimetinib (S8041) and MK-2206 (S1078) (Selleckchem).

### Maintenance cell culture

The human hepatoma cell lines: HepG2 IGF1R KO, HepG2 WT (ATCC-HB8065), and Hep3B (ATCC-HB8064) were cultured in DMEM, 4.5 g/L D-glucose (Gibco). The human breast cancer cell lines; BT549, BT474, HCC38, and HCC1937 were cultured in RPMI 1640 (Gibco), all supplemented with 10% fetal bovine serum (FBS) and penicillin (100 U/mL) and streptomycin (100 μg/mL) (P/S) (Gibco). The rat hepatoma cell line, H4IIE (ACTT-CRL-1548), was cultured in MEM media (Gibco) with 1% MEM Non-Essential Amino Acids Solution (Gibco) and 1% pyruvate (Gibco) and supplemented with 10% FBS and P/S. The human AML cell lines MOLM-13 (ACC-554), TPH1 (TIB-202), and EOL-1 were cultured in RPMI with Glutamax (Gibco), supplemented with 10% FBS and P/S. Additionally, for the TPH1 cells, 20 nM 4-(2-hydroxyethyl)-1-piperazineethanesulfonic acid (HEPES) (Gibco) and 50 µM 2-Mercaptoethanol (Gibco) were added, while 20 nM glutamine (Gibco) were added for the EOL-1 cells. The HepG2 IGF1R KO and HepG2 WT cells were cultivated on collagen-coated surfaces.

### IGF1R KO in HepG2

The HepG2 IGF1R KO cell line was generated by transfecting HepG2 WT with Cas9 protein, crRNA targeting exon1, and tracrRNA in IDTE buffer (Integrated DNA Technologies) using Neon transfection kit (Invitrogen). The transfected cell pool was expanded from 24W plate to T75 flask. Using human IGF1R PE-conjugated antibody (R&D systems), the IGF1R negatively stained population was bulk-sorted using SH800 cell sorter (Sony) to select for IGF1R KO cells. After expansion, the cell pool was limited diluted into Nunclon delta surface 96W plates (Thermo Scientific) using medium supplemented with 50% conditioned medium. Surviving clones were expanded in poly-D-lysine coated plates (Corning) and screened with Zero Blunt TOPO PCR cloning (Invitrogen) and Sanger sequencing (Eurofins), leading to the selection of the HepG2 IGF1R KO clone.

### Cell treatments

To mimic insulin sensitivity and resistance in HepG2 IGF1R KO cells, cells were seeded in DMEM 1 g/L D-glucose (Gibco) with 10% FBS and 1% P/S. The following day media was exchanged to DMEM, 1 g/L D-glucose with 1% P/S for 48 hours (serum washout). Next, media was exchanged to DMEM, 1 g/L D-glucose with 1.25% human serum albumin (HSA) (Sigma) and either physiologic (0.1 nM) or pathologic (3 nM) insulin for 48 hours. The insulin-sensitivity and - resistance protocol was adapted from Dall’Agnese *et al*. [17].

For experiments on the cell line panel, cells were seeded in their respective growth media and serum-starved overnight in media without FBS. After serum washout, cells were either continued serum-starved or treated with 3 or 100 nM insulin with 1.25% HSA for 24 hours.

In the ERK and AKT inhibition experiment, HepG2 IGF1R KO cells were serum-starved overnight in DMEM 1g/L D-glucose with 1% P/S. Next pretreated with cobimetinib (MEK inhibitor) or MK-2206 (AKT inhibitor) for 30 minutes, followed by 24 hours co-treatment with 3 nM insulin for 24 hours.

### Cell stimulation and lysis

To study IR activation in insulin-sensitive and -resistant HepG2 IGF1R KO cells, an extensive seven-step insulin wash-out was performed with DMEM, 1 g/L D-glucose media. The procedure comprised three continual media exchanges followed by three 5-minute exchanges, and one 20-minute exchange, with incubation in a cell incubator for the 5- and 20-minute steps.

After the insulin wash-out, cells were stimulated for 5 minutes with either 0, 0.1, 3, or 100 nM insulin in DMEM, 1 g/L D-glucose with 1.25% HSA in a cell incubator. Cells were washed and lysed directly in experiments without specified insulin stimulation. Wash was performed with phosphate-buffered saline (PBS) and lysis with co-IP lysis buffer [50 mM tris-HCl (pH 7.5), 150 mM NaCl, 1 mM calcium chloride, 1% Triton X-100] added 5 mM β-glycerophosphate, 5 mM sodium fluoride, 1 mM sodium ortho-vanadate, and one cOmplete EDTA-free Protease inhibitor tablet (Roche) per 10 mL solution. For SDS-proteome analysis, lysates were collected after a 48-hour differential insulin treatment with 0.1 nM (insulin-sensitive condition) or 3 nM insulin (insulin-resistant condition), without 5-minute insulin stimulation. Cell pellets were washed twice with PBS and lysed by heating at 95°C for 10 minutes in lysis buffer consisting of 100 mM Tris-HCl (pH 8.5), 5% SDS, 5 mM TCEP, and 10 mM CAA. Sonication was performed thereafter. For all samples protein concentration was determined using BCA Protein Assay Kit (Pierce).

For the quantitative SILAC MS-based proteomics and phosphoproteomics experiments, H4IIE cells were subjected to SILAC for a minimum of 14 days. The SILAC media was composed of MEM for SILAC (Thermo Scientific), 1% P/S, 10% dialyzed FBS for SILAC (Gibco), 1% MEM Non-Essential Amino Acids Solution, sodium pyruvate, and GlutaMAX (Gibco). Three different cell populations were achieved by the addition of 166 µM lysine (Lys) and 274 µM arginine (Arg) in the following variations: Light: L-Arg0 and L-Lys0 (Sigma) Medium: Arg6 (L-(13C6) Arg) and Lys4 (L-(2H4) Lys). Heavy: Arg10 (L-(13C6, 15N4) Arg) and Lys8 (L-(13C6, 15N2) Lys). The medium and heavy labeled amino acids were obtained from Cambridge Isotope Laboratories (Tewksbury).

For phosphoproteome analysis, SILAC H4IIE cells were serum-starved overnight in SILAC media containing 0.1% FBS, before 100 nM insulin, 100 nM S597, or vehicle was added for 5, 15, or 30 minutes. At the end of stimulation, cells were washed five times in ice-cold DPBS and lysed in 6 M guanidine hydrochloride (Sigma-Aldrich) in 10 mM Tris, pH 8, added PhosSTOP (Roche). Protein concentrations were measured by BCA. For one biological replicate, 2 mg protein from each treatment condition was mixed 1:1:1, to give a final protein content of 6 mg and for the second replicate, a 1:1:1 mixture was composed of 4 mg from each condition.

For proteome analysis, SILAC H4IIE cells were treated for 24 or 48 hours with regular SILAC media containing 100 nM human insulin, 100 nM S597, or vehicle. For the-48 hour stimulation, the media was changed after 24 hours of stimulation. Following treatment, the cells were washed five times in ice-cold DPBS and lysed with 0.5% RapiGest (Waters) in 50 mM triethylammonium bicarbonate (TEAB) buffer. Cell lysates were collected and 1 µL Benzonase nuclease (EMD Chemicals) was added. Protein concentrations were measured by BCA. Finally, 200 µg proteins from each treatment condition were mixed giving a 1:1:1 protein mixture ready for digestion.

### Cell viability assay

HepG2 IGF1R KO cells were seeded in black/clear bottom 96-well cell culture plates (Fisher Scientific) and subjected to the insulin resistance induction protocol. As described, after 48-hour serum-starvation, cells were cultured with 0.1 or 3 nM insulin in DMEM, 1 g/L D-glucose with 1.25% HSA for 48 hours. Cell viability was assessed using the CellTiter-Glo® Luminescent Cell Viability Assay (Promega) following the manufacturer’s instructions. Luminescence was measured at 37°C using a CLARIOstar® plate reader.

### Assay for insulin receptor and AKT activation

Cells were seeded in 96-well cell culture microplates (Fisher Scientific) and subjected to the insulin-resistant protocol as described in the previous section. A seven-step insulin washed-out was performed before 5-minute insulin stimulation with doses ranging from 0 to 600 nM. Reagents were supplied in the AlphaScreen, SureFire, Insulin Receptor (p-Tyr1150/1151), and AKT 1/2/3 (p-S473) Assay Kits (PerkinElmer). Cell lysis and SureFire Assay were performed following the manufacturer’s instructions.

### siRNA experiments

HepG2 IGF1R KO cells were transfected in 6-well cell culture plates (Fisher Scientific) using Lipofectamine RNAiMAX Transfection Reagent (Thermo Fisher Scientific). Following 48-hour serum-free incubation, the cells were treated with the transfection reagent in DMEM, 1 g/L D-glucose, supplemented with 1.25% HSA and either 0.1 nM or 3 nM insulin for 12 hours. The EphA2 siRNA SMARTpool ID: L-003116-00-0005 (Dharmacon Inc.) or the ON-TARGETplus Non-targeting Control SMARTpool ID: D-001810-10-05 (Dharmacon Inc.) was added to a final concentration of 48 nM. After 12 hours of transfection, the medium was exchanged and the cells were cultured with insulin for an additional 36 hours, for 48 hours of incubation, following the insulin resistance-inducing protocol. Subsequently, the insulin was washed-out, and the cells were stimulated with insulin as previously specified before being lysed for WB analysis.

### Western blotting

SDS-PAGE (4-12% Bis-Tris, MOPS buffer, NuPAGE, Invitrogen) was blotted to nitrocellulose transfer membranes according to the manufacturer’s protocol (iBlot). SuperSignal West Dura Extended Duration Substrate or West Pico Chemiluminescent Substrate (Thermo Scientific) was used for detection with an LAS-3000 Imaging System (Fuji). Reprobing of blots after restoring Western Blot Stripping Buffer (Thermo Scientific). Quantitative analysis of bands of interest was done using either Image Gauge v.4.0 or ImageJ software tools.

### Flow cytometry

Cell surface IR receptor levels were quantified using the QIFIKIT kit (DAKO) for flow cytometry analysis. HepG2 IGF1R KO cells were cultured in 6-well plates and detached with Versene (Gibco) for 5 minutes at 37°C. The cells were washed and resuspended in cold FBS Stain Buffer (BD Biosciences). 2×10^4^ cells per condition were stained with anti-IR (D2) mouse IgG1 or isotype control and incubated in the dark for 1 hour at 4°C followed by three washing steps in cold FBS Stain Buffer. Subsequently, the cells were incubated at room temperature in the dark for 45 minutes with anti-mouse secondary antibodies conjugated to Alexa Fluor (FITC). After three washing steps, the cells were resuspended in 200 µL FBS Stain Buffer for flow cytometric analysis.

Flow cytometry data was acquired using a NovoCyte Quanteon flow cytometer, recording 15,000 events of the desired and gated populations. The instrument was operated with a sample volume of 75-100 µL, a flow rate of 35 µL/sec, and the FITC channel set at 480 nm for fluorescence detection. Data analysis was performed using NovoExpress v.1.5.6 software, and single-cell populations were gated based on a plot of side scatter (SSC-A) versus forward scatter (FSC-A) light signals.

### Real-time PCR and RNA sequencing

RNA was extracted after 24-hour culturing in DMEM, 1 g/L D-glucose, 10% FBS, with 1% P/S, after 48 hours in DMEM, 1 g/L D-glucose with 1% P/S, and after inducing insulin sensitivity and resistance according to the protocol described above. RNA extraction was performed using the RNAdvance Cell v2 Kit (Beckman Coulter). The RNA concentration was determined using NanoDrop™ 8000, and 300 ng was used as starting material for cDNA synthesis, applying the iScript cDNA Synthesis Kit (Bio-Rad). Real-time PCR was performed on a Thermo Fisher Scientific QuantStudio 12K Flex using TaqMan™ Fast Advanced Master Mix (Thermo Fisher). TaqMan primers (Thermo Fisher) for the IR, EphA2, and PPIB genes were used with the following IDs: INSR: Hs00961554_m1, EPHA2: Hs01072272_m1, PPIB: Hs00168719_m1. For each condition, the average of 12 technical replicates was used as the representative Ct value of each biological replicate (n=4). The Ct values were normalized by determining the expression levels of the target genes relative to the endogenous housekeeping gene PPIB. To correct the expression of the target gene relative to PPIB, the ΔCt method was applied, where ΔCt is the difference in Ct values between the target gene and PPIB. The relative quantification (RQ) was calculated using the formula RQ = 2^-ΔCt^ [35]. The corrected individual Ct values were normalized to the average of the biological replicates for the chosen reference condition.

### Immunofluorescence imaging

Immunofluorescence staining was performed on HepG2 IGF1R KO cells, which were rendered insulin-sensitive or insulin-resistant and subsequently stimulated with 3 mM insulin for 5 minutes as previously described. The cells were fixed in 4% paraformaldehyde in PBS for 10 minutes at room temperature. After fixation, the samples were washed with PBS and permeabilized by incubation with 0.5% Triton X-100 in PBS for 10 minutes at room temperature. Following permeabilization, the samples were washed again with PBS to remove residual detergent and then blocked in 3% bovine serum albumin (BSA) (Sigma) in PBS for 1 hour at room temperature. Plates were incubated overnight at 4°C with the appropriate primary antibodies. The primary antibodies were diluted in 2% BSA in PBS, with a dilution of 1:800 for the 83-7 IR antibody overnight. The samples were washed three times with PBS. Subsequently, the samples were incubated with the appropriate fluorescently labeled secondary antibodies in 3% BSA at room temperature for 1 hour in the dark. The samples were washed in PBS and incubated with Hoechst dye diluted 1:5000 in PBS for 2-5 minutes at room temperature. The samples were washed with PBS before imaging using the Opera Phenix® Plus Imager (PerkinElmer) with a ×63 objective. Either 7 or 18 stacks were acquired, covering a total depth of 3 μm. The acquired images were processed, and scale bars were determined using Harmony v.5.1 software (PerkinElmer). Quantitative values corresponding to the fluorescence signal intensity in the regions containing the cells were determined for each image in the dataset. The same settings and parameters were consistently applied to all images.

### Sample preparation for mass spectrometry-based proteomics

For co-IP with the IR for MS-based IR interactome analysis, the IR-specific 83-7 antibody was coupled to Dynabeads™ MyOne™ Tosylactivated magnetic beads (Invitrogen™) following the manufacturer’s instructions. Non-conjugated beads were prepared in parallel, with no addition of antibody in the conjugation step. 500 µg of protein was used for IR co-IP. The lysate was diluted to a concentration of 0.5 mg/mL in IP-bead incubation buffer [50 mM HEPES; 125 mM NaCl; 0.5 mM CaCl_2_; 5 mM MgSO_4_; 0.0125% Tween20; 0.5% TritonX100] and 100 µg of non-conjugated beads were added to each sample for pre-clearing. This was done in a 96-deep well format, with samples incubated shaking at 750 rpm at 4°C for 1 hour. While keeping samples cold, the supernatant was carefully transferred to a new 96-well plate to prevent any bead contamination. Afterward, 150 µg of antibody-conjugated beads were added to each sample, followed by incubation while shaking at 750 rpm at 4°C for 4 hours. After the IR co-IP, the beads were passed through an automated washing procedure on a KingFisher robot with five 1-minute washing steps in the IP-bead incubation buffer. Finally, the protein was eluted using 100 µL of IgG Elution buffer (Pierce™).

For MS-based single-shot proteome and phosphoproteome analysis, each sample was digested and then split for either direct single-shot proteome or phosphopeptide enrichment. A total of 500 μg of protein from the co-IP lysate and the SDS-lysate were separately digested overnight at 37°C on a Kingfisher Flex robot using the protein aggregation capture (PAC) method [36], [37]. The digestion was performed in a 50 mM TEAB buffer with 0.2 μg of Lys-C (Wako Chemicals) and 0.4 μg of trypsin (Promega). Trifluoroacetic acid (TFA) acidification was applied to quench the digestion, followed by loading of 750 ng of peptides onto Evotip Pure (Evosep) disposable trap columns for single-shot proteome analysis of both lysates. The remaining peptides from the digested co-IP lysate, corresponding to 200 μg peptide mixtures, underwent clean-up using SepPak (C_18_ Classic Cartridge, Waters). Elution was performed using 40% and 60% acetonitrile (ACN). The samples were then lyophilized, resuspended in a loading buffer composed of 80% ACN, 5% TFA, and 1 M glycolic acid, and subjected to automated Ti-IMAC (titanium immobilized metal affinity chromatography) phosphopeptide enrichment [37], [38]. TiIMAC-HP beads (MagReSyn, Resyn Biosciences) were incubated with the peptides for 20 minutes, followed by successive washing steps with the loading buffer, 80% ACN, 1% TFA, and 10% ACN, 0.2% TFA. Phosphopeptides were eluted in 1% ammonia and clarified by filtration using a 0.45 μm filter plate (Millipore, Sigma–Aldrich) before being loaded onto Evotip Pure trap columns.

For SILAC H4IIE phosphoproteome, reduction and alkylation were performed by the addition of 5 mM TCEP (Sigma-Aldrich) and 4.5 mM CAA diluted in 100 mM TEAB buffer. Samples were digested with Lys-C (1:100) at room temperature for 4 hours and subsequently diluted in 10 mM Tris, pH 8, to a final guanidine hydrochloride concentration of 1.5 M. Next, digestion with sequencing grade modified trypsin 1:100, was performed overnight at 37°C. Samples were acidified with TFA and centrifuged at 3000 g for 10 minutes. The peptide mixtures were desalted and cleaned using SepPak with the final wash was done in MilliQ H2O and peptides were eluted in 50% ACN. The eluted peptides were lyophilized before phosphopeptide enrichment.

pTyr IP was performed using 120 µL Tyr1000 and 40 µL pTyr100 Ab combined per pull-down. Lyophilized samples were reconstituted in 1.4 mL of IP buffer provided with the kit and performed according to the manufacturer’s protocol. The pTyr enriched samples were purified on a homemade C8 STAGE-Tip and analyzed by LC-MS/MS. The remaining sample was high-pH (HpH) fractionated followed by titanium dioxide (TiO2) enrichment performed as previously described [39]. In short, samples were eluted from the Sep-Pak and lyophilized to a remaining volume of approximately 1 mL. The samples were fractionated using a Waters XBridge C18 3.5 µm, 4.6 x 250 mm column on an Agilent 1290 Infinity HPLC system connected to a Rheodyne MX Series II valve for sample injection. The system was operating at 1 mL/minute. Buffer A was 10 mM ammonium hydroxide and buffer B was 90% ACN and 10 mM ammonium hydroxide. Samples were loaded into the column at 1 mL/minute for 15 minutes in 1% buffer B followed by the gradient: (1-25% B in 50 minutes, 25-60% B in 4 minutes, 60-70% B in 2 minutes 70% B in 5 minutes, and 1% B for 4 minutes)

The HpH fractions were lyophilized and combined into 12 fractions. Phosphopeptides were enriched using metal oxide affinity enrichment (MOAC) with 5 µm Titansphere TiO2 beads (GL Sciences, Japan) as described elsewhere [40], [41]. The TiO2 beads (3.5 mg per 1 mg sample) were incubated in 2,5-dihydroxybenzoic acid (DHB) (0.02 g/mL in 80% ACN, 6% TFA) for 20 minutes. The samples were lyophilized and reconstituted in 1 mL 80% ACN, 6% TFA and next incubated with freshly prepared TiO2 in DHB for 30 minutes at room temperature. The supernatant was collected for each fraction and pooled into three samples for a second round of TiO2 enrichment, which afterward was pooled into a single sample and subjected to a third round of TiO2 enrichment. The TiO2 beat pellets from the 16 TiO2 enriched samples were washed on a C8 STAGE-tip with 10, 40, and 80% ACN in 6% TFA, respectively. Next, the enriched phosphopeptides were eluted with 5% ammonium hydroxide, followed by 10% ammonium hydroxide with 25% ACN. The samples were lyophilized to a remaining 4 µL and 20 µL 5% ACN, 1% TFA was added to each sample, and the samples were purified on C18 STAGE-tips before the LC-MS/MS analysis.

For SILAC proteome samples were reduced using 5 mM dithiothreitol (DTT) and heating of the samples to 60°C for 30 minutes and subsequent alkylation with 10 mM CAA for 30 minutes at room temperature. Samples were incubated with Lys-C (1:360 w/w) at 30°C for 3 hours followed by incubation with trypsin (1:100, w/w) at 37°C overnight. To remove the RapiGest, TFA was added to a final concentration of 0.5% and heated to 37°C for 30 minutes and the supernatant was collected after centrifuging for 10 minutes at 13000 rpm. Samples were cleaned on Sep-Pak columns before fractionation. HILIC fractionation of the peptides was performed using a TSK-Gel amide-80 3UM 2 × 150 mm SILICA HPLC column (Tosoh Bioscience) connected to a 1290 Infinity Binary LC system from Agilent Technologies (Santa Clara). The system was operating at 0.2 mL/minute. Buffer A (0.1% formic acid (FA)) and buffer B (98% ACN with 0.1% FA). Approximately 200 µg digested protein was cleaned by SepPac, lyophilized, and reconstituted in 4 µl buffer A and 36 µl buffer B by mixing at 1100 rpm for 30 minutes. The samples were loaded onto an HPLC column at a flow rate of 0.2 mL/minute for 5 minutes using 5% buffer A. The gradient progressed from 5-100% A for 47 minutes, 100-5% A for 1 minute, and 5% A for 15 minutes. Fractions were pooled into 10 fractions, lyophilized, and reconstituted in 12 µL 0.1% FA and 1% TFA for 30 minutes mixing at 1200 rpm. 5 µL of each sample were then analyzed by LC-MS/MS.

### Liquid chromatography-tandem mass spectrometry and data analysis

All HepG2 IGF1R KO proteomics samples were analyzed using the Evosep One system, coupled with the Orbitrap Exploris 480 MS (Thermo Fisher Scientific) using Xcalibur tune v.1.1. A 15 cm, 150 μm inner diameter capillary column packed in-house with 1.9 μm Reprosil-Pur C18 beads (Dr. Maisch) was used. The analysis was done using pre-programmed gradients of 30 samples per day (30SPD) for the interactome and single-shot proteome, and 60 samples per day (60SPD) for phosphoproteome analysis. The column temperature was set to 60°C using the integrated PRSO-V1 column oven (Sonation).

The spray voltage was set to 2 kV, the funnel RF level was 40, and the heated capillary temperature was 275°C. All analyses were acquired in data-independent acquisition (DIA) mode, full MS resolution was set to 120,000 at m/z 200. The full MS AGC target was 300% with an injection time (IT) of 45 ms, and the mass range was 350-1400 m/z. The AGC target value for fragment spectra was 100%. In DIA analysis, 49 windows of 13.7 m/z were scanned from 361 to 1033 m/z, with a 1 Da overlap. The resolution for these scans was 15,000, with an IT of 22 ms, and the normalized collision energy was 27%. For phosphoproteome analysis using DIA, 17 windows of 39.5 m/z were scanned from 472 to 1143 m/z, with a 1 m/z overlap. The resolution for these scans was 45,000, with an IT of 86 ms, and the normalized collision energy was 27%. All data were acquired in positive ion and profile mode.

The acquired DIA raw data were analyzed using Spectronaut v.17.0 with a library-free approach (directDIA) [42]. The analysis utilized the SwissProt human database, UP000005640_9606 release 2018 with signal peptides removed and a separate database of common contaminants. For interactome and proteome analysis, fixed modifications included carbamidomethylation of cysteine, while protein N-terminal acetylation and oxidation of methionine were set as variable modifications. For the interactome searches, cross-run normalization was disabled and the protein LFQ method was changed to QUANT 2.0. For phosphoproteome analysis, phosphorylation of serine, threonine, and tyrosine were included as variable modifications.

All H4IIE SILAC samples were analyzed on an Easy-nLC 1000 connected to a Q-Exactive Orbitrap Plus (Thermo Scientific). The peptides were separated on C18-bead columns packed in-house. Analyzed with a 160 minutes gradient for the proteome analyses (4-25% buffer B in 120 minutes, 25-40% buffer B in 20 minutes, 40-80% buffer B in 2 minutes), a 240 minutes gradient for the pTyr samples (4-25% buffer B in 190 minutes, 25-40% buffer B in 30 minutes, 40-80% buffer B in 2 minutes, and a 165 minutes gradient for the TiO_2_ enriched phospho-samples (4-25% buffer B in 120 minutes, 25-40% buffer B in 25 minutes, 40-80% buffer B in 2 minutes).

The Q-Exactive mass spectrometer was operated in positive ion mode with a capillary temperature of 275°C. The data were acquired in data-dependent acquisition (DDA) mode with a loop count of 12. The resolution for the full scan was 70,000, AGC target of 3E6, maximum IT 20 ms, and 1 microscan. The MS/MS scans for the phospho-samples were recorded with a resolution of 35,000, AGC target of 1E6, maximum IT 120 ms, isolation window 2.2 m/z, NCE of 25, and underfill ratio 1.0% (intensity threshold of 8.3E4). The dynamic exclusion was set to 20 seconds. For the proteome samples, the MS/MS scans were recorded with a resolution of 17,500, AGC target of 5E5, maximum IT 50 ms, isolation window 2.2 m/z, NCE of 25, and underfill ratio 1.0% (intensity threshold of 1E5). The dynamic exclusion was set to 20 seconds.

Raw MS files from the H4IIE SILAC phosphoproteome and proteome analysis were processed using the MaxQuant software [43] version 2.1.4.0. The precursor MS signal intensities were determined and the SILAC triplets were automatically quantified. Proteins were identified by searching the complete rat UniProt database, UP000002494_10116 release 2022. MS/MS spectra were matched with a mass tolerance of 20 ppm. The proteome was searched with carbamidomethyl of cysteine as a fixed modification and oxidation of methionine and acetyl protein N-term as variable modifications. The peptide and protein false discovery rate (FDR) was set at 1%. For the phospho-samples, phosphorylation of serine, threonine, and tyrosine was included as variable modifications, and PSM FDR was set at 1%. Match between runs, with a window of 0.7 minutes, and a maximum of 2 missed cleavages were activated for proteome and phosphoproteome data.

### Bioinformatics analysis of proteomics data

All datasets were initially filtered to remove proteins identified as contaminants from the common contaminant database before downstream analysis.

For the IR interactome, all files were searched in Spectronaut. Subsequently, the data was log-transformed, filtered, normalized, and imputed using ProStaR v.1.30.7 online software tool [44]. Samples were grouped and filtered to include protein IDs identified in at least 4 out of 5 samples for at least one condition (Table S1). Normalization was performed using the global quantile alignment method. Imputation was carried out in two steps using the structured least square adaptive method for partially observed values (POVs) and DetQuantile for missing values on entire conditions (MEC) partially observed values and DetQuantile for missing values on an entire condition. Proteins annotated in the CRAPome database v.2.0, with a score above 3 based on 718 datasets, were excluded [45]. Based on DeepLoc prediction of subcellular localization, extracellular and nuclear proteins were excluded [46].

For the dynamic interactome, data were processed in ProStaR as described above, however with insulin-sensitive and insulin-resistant samples processed individually to account for uneven efficiency. Volcano plots were generated to visualize the differential recruitment of interactors upon insulin stimulation with different concentrations. The plots were created by plotting the - log10-transformed p-values derived from a two-sided t-test against log2-transformed fold changes. Statistical significance was determined based on a hyperbolic curve threshold with s0=0.1 and FDR<0.05, derived from statistical analysis using the Perseus v.1.6.5.0 software [47] (Table S1).

For the phosphoproteomics and single-shot proteomes, cross-run normalization was performed in the Spectronaut software. For the phosphoproteomics dataset, phosphorylation sites with a localization probability score of ≥0.85 were log-transformed using ProStaR, followed by grouping into conditions and filtering for protein identification in minimum 3 out of 5 samples for at least one condition (Table S2). To account for the intrinsic complexity of DIA spectra and increase confidence in the site localization of phosphorylation sites, the site probability cut-off was set to 0.85. Imputation was conducted using SLSA for POVs and DetQuantile for MEC. Median subtraction was performed, based on biological experiments before the generation of volcano plots to illustrate the differential regulation of phosphorylation sites upon insulin stimulation with different concentrations for insulin-sensitive and insulin-resistant samples and between insulin-sensitive and -resistant cells directly. Statistical significance was defined such that phosphorylation sites were found to be significantly regulated when the change was minimum 2-fold and p-value<0.05, or as specified in the respective figure legends.

The single-shot proteome was log2-transformed using the Perseus software, and median subtraction was performed, based on biological experiment. Volcano plots were generated as described above. The 5-minute insulin stimulation was not considered in the analysis of differentially regulated proteins. For the co-IP lysis buffer proteome, the cutoffs for both directions were set to a ratio cutoff of 1.25 and FDR<0.05. For the SDS-based proteome, statistical significance was determined based on a hyperbolic curve threshold with s0=0.1 and FDR<0.05 derived from statistical analysis using Perseus (Table S3). The protein association network based on IR interactome data was obtained using the STRING database v.11.5 [48]. Default search parameters were used in the multiple protein tab. The network was visualized using Cytoscape v.3.10.0 [49]. The KEGG pathway enrichment analyses were performed using the InnateDB resource v.5.4 [50]. The pathway overrepresentation analysis was conducted using the hypergeometric algorithm, with Benjamini-Hochberg FDR correction (Ben. Ho. FDR corrected). Enriched pathways were defined with pathway ORA p-value<0.05. If multiple enriched pathways had an identical profile of identified involved proteins, only the entry with the lowest pathway ORA p-value was included.

The H4IIE cell SILAC proteome and phosphoproteome datasets were filtered for contaminants and reverse hits. For the proteome analysis with 24- and 48-hour insulin and S597, treatment ratios and intensities were log2-transformed. Normalized ratios and intensities were reported as the median of three replicates. Proteins identified by only one peptide were excluded, and data were further filtered to include proteins with at least two valid values (of the three replicates) in at least one of the treatment groups. Statistically significant changes in protein ratio induced by long-term treatment with insulin or S597 were determined using significance B testing (p < 0.05) with the Perseus software (Table S4).

For the DDA SILAC phosphoproteomics dataset, only phosphorylation sites defined based on a phosphorylation site localization probability ≥0.75 (class I, as defined by Olsen *et al*. [51]) were included. Normalized SILAC phosphorylation site ratios and intensities were reported as the median of two replicates, and the data was log2-transformed. The table was expanded to include singly, doubly, and triply phosphorylated sites. Only relative changes compared to untreated controls were used; therefore, H/M SILAC ratios were not included in the downstream analysis. A cutoff of >2-fold regulation, if valid in at least one of the treatment groups, was applied as section criteria (Table S4).

### Proteomics data availability

The DIA and DDA MS proteomics data have been deposited to the ProteomeXchange Consortium via the PRoteomics IDEntifications (PRIDE) [52] partner repository with the dataset identifier PXD047011 and PXD046865, respectively.

## Results

### Insulin resistance in a hepatocyte cell line model leads to decreased levels of insulin receptor and AKT phosphorylation

To investigate the mechanisms of insulin resistance in a hepatocellular context, we employed HepG2 cells with a CRISPR IGF1R KO to mimic hepatocytes of the liver, known to lack IGF1R expression [32], [53]. A cell model system of in situ hepatocyte insulin sensitivity and resistance was established using a protocol described in Dall’Agnese *et al.* [17]. In the applied model, HepG2 IGF1R KO cells were cultured for 48 hours in serum-starvation media, followed by a 48-hour stimulation with concentrations of 0.1 nM (physiological) or 3 nM (pathological) insulin to generate insulin-sensitive and -resistant cells, respectively (Figure S1.A).

HepG2 IGF1R KO cells exposed to a pathological insulin concentration exhibited hallmark characteristics of insulin resistance, such as decreased levels of AKT phosphorylation upon 5-minute insulin stimulation (Figure 1.A-B). Moreover, approximately 30% decreased level of IR in insulin-resistant cells was observed compared to insulin-sensitive cells (Figure 1.A and C, and Figure S1.B). This finding is consistent with other studies showing decreased IR levels in insulin-resistant states [13], [14], [54]–[56]. Decreased *INSR* mRNA levels were shown in the insulin-resistant cells, compared to the -sensitive (Figure S1.C). Flow cytometry showed that the cell surface IR levels reflected the decrease in total IR protein level in the insulin-resistant cells (Figure 1.D-E). Immunofluorescence imaging further supported the higher level of IR in the insulin-sensitive cells and that the subcellular localization of IR varied between insulin-sensitive and - resistant cells (Figure 1.F). The insulin-resistant cells showed approximately 33% enhanced cell viability compared to the -sensitive cells (Figure S1.D), despite their impaired insulin response involving AKT.

**Figure 1.**
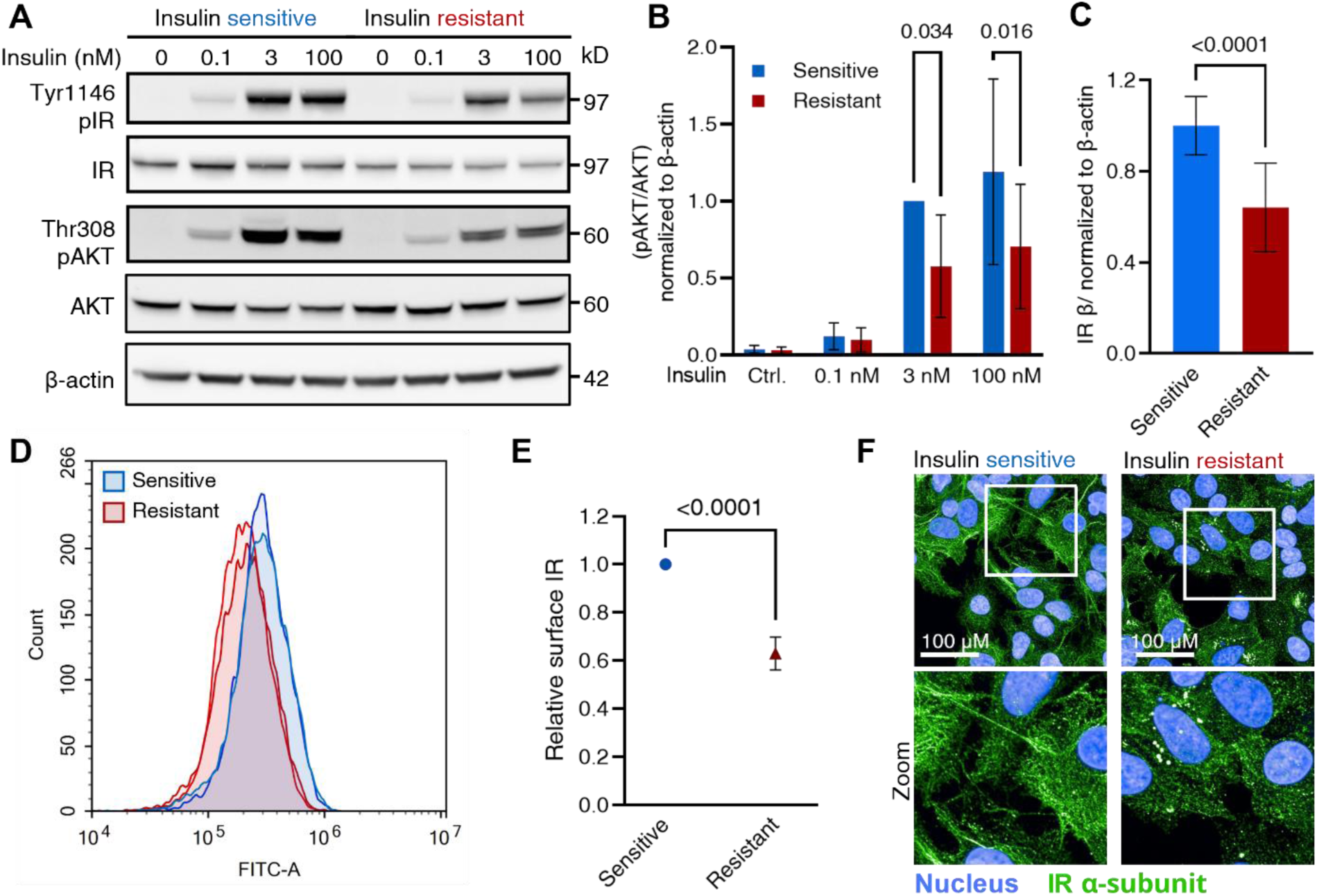
Reduced IR protein and AKT phosphorylation levels in insulin-resistant HepG2 IGF1R KO cells. **A**. Immunoblot of insulin-sensitive and -resistant HepG2 IGF1R KO cell lysates. The cells were stimulated with increasing insulin concentrations for 5 minutes (representative blot of n=4 independent experiments). **B**. Phospho-AKT levels quantified from the immunoblot shown in A and normalized to total AKT and β-actin. Insulin-sensitive and -resistant cells were stimulated with 0.1, 3, and 100 nM insulin. Unstimulated control included. Data presented as means ±SD of n=3 independent experiments relative to insulin-sensitive cells stimulated with 3 nM insulin. p-values<0.05 are indicated (2-way ANOVA). **C**. Quantification of IR levels, relative to β-actin under insulin-sensitive and -resistant conditions, independent of 5-minute insulin stimulation. The data are represented as means ±SD relative to the insulin-sensitive cells (two-sample unpaired t-test) (four technical replicates from n=4 independent experiments). **D**. Flow cytometry histograms showing the average FITC signal for cell-surface IR in insulin-sensitive (blue) and -resistant (red) HepG2 IGF1R KO cells. Representative of n=4 independent experiments in technical duplicates. **E**. Quantification of surface IR levels in insulin-resistant relative to -sensitive cells, based on flow cytometry data in D. **F**. Representative immunofluorescence images of IR (green) with nuclear staining (blue) in insulin-sensitive and insulin-resistant HepG2 IGF1R KO cells. The lower panel shows a zoomed-in view.

These initial experiments confirmed that the HepG2 IGF1R KO cell model displayed the expected traits of insulin resistance, thus making it a suitable model for the analyses of hepatocellular insulin resistance.

### Multi-layered proteomics datasets characterize the molecular dynamics of insulin signaling in insulin sensitivity and resistance

The early signaling effects of the insulin dose-response were systematically investigated by multi-layered proteomics analysis including, the interactome, to identify proteins involved in IR proximal signaling events; and the phosphoproteome, studying the associated signaling cascades in the insulin-resistant HepG2 IGF1R KO cell model (Figure 2.A-B and Figure S2.A). To study the IR interactome, a co-IP of the IR was performed, followed by label-free DIA liquid chromatography-tandem mass spectrometry (LC-MS/MS) analysis. A total of 5,563 proteins were identified with good correlation between samples (Pearson correlation in the range of 0.63-0.95) (Figure S3.A and Table S1). To ensure the quality of IR enrichment in our endogenous IR-expressing HepG2 IGF1R KO cell model, IR enrichment was confirmed by immunoblot (Figure S3.B). Additionally, the IR was identified with the highest number of assigned peptide precursors (202) and a sequence coverage of 73% in our MS interactome dataset (Figure 2.C and Table S1). These quality checks together confirmed the generation of a dataset enabling analysis of IR signaling in a cell model with endogenous IR levels. Furthermore, the downregulated IR levels in insulin-resistant compared to -sensitive cells were reflected in the IR levels after enrichment for the two unstimulated conditions (Figure S3.C). Affinity purification-MS (AP-MS) analyses involve the identification of proteins that interact with a specific target protein, and these analyses are often challenged by a high degree of nonspecific binding. To distinguish true IR interactors from unspecific background binders, the dataset was refined using the CRAPome (Contaminant Repository for Affinity Purification), a published repository of commonly detected contaminating proteins in AP-MS studies [45]. Applying the CRAPome enhanced the accuracy of the list of potential IR binding partners and helped distinguish true binding candidates from background noise. After this, 1,664 proteins remained, and among these, 207 proteins showed a ≥10-fold increase in the number of peptide precursors compared to proteome background levels. This increase indicates an enrichment within the IR co-IP, highlighting proteins likely specific to IR proximate signaling. The ≥10-fold criterion enhances the stringency of the analysis by prioritizing substantial changes and reducing false positives (Figure 2.D, Figure S3.D-E, and Table S1).

**Figure 2.**
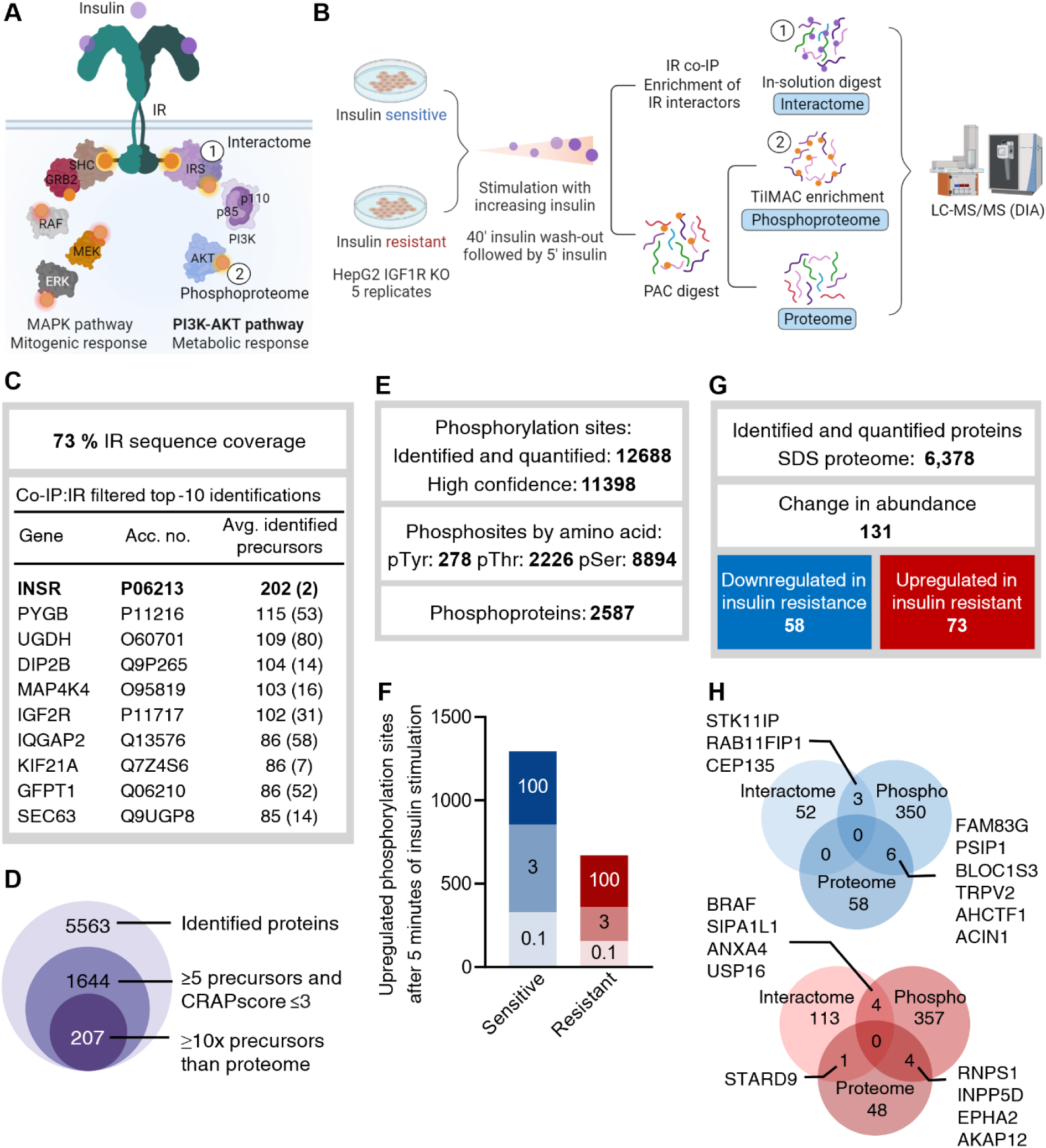
Comprehensive multi-layered proteomics analysis depicts global signaling changes in induced insulin resistance. **A**. Schematic illustration of canonical insulin signaling pathways through the IR. **B**. Workflow illustration for DIA MS-based interactome, phosphoproteome, and single-shot proteome analyses. Cell lysates of insulin-sensitive and -resistant cells stimulated with 0.1, 3, and 100 nM insulin for 5 minutes were collected. See Figure S2.A. **C**. IR sequence coverage after co-IP and listing of top 10 proteins in the co-IP dataset sorted by MS-based identified precursors, after filtration. **D**. Number of proteins in the IR co-IP dataset after analysis filtering steps, including filtering using the Contaminant Repository for Affinity Purification (CRAPome). **E**. Phosphoproteome summary: Number of identified phosphorylation sites and phosphoproteins. **F**. Number of phosphorylation sites with upregulated levels in the phosphoproteome after insulin stimulation in insulin-sensitive and -resistant cells. Bar colored based on insulin concentration used for 5-minute stimulation. **G**. Overview of proteome data: Proteins identified from the SDS-proteome dataset specifying regulated proteins in insulin-sensitive and -resistant cells. Significance based on volcano-plot in Figure 5.A made in Perseus (two-sided t-test, FDR<0.05, s0=0.1). **H**. Venn diagram showing overlap of upregulated proteins across the three layers of proteomics analysis. Insulin-sensitive and -resistant cells are shown in blue and red, respectively.

To characterize insulin-stimulated global phosphorylation site changes in insulin-sensitive and -resistant cells, a quantitative phosphoproteomics experiment was performed (Figure 2.B), based on the enrichment of phosphorylated peptides using magnetic titanium-chelated metal ion affinity chromatography (Ti-IMAC) beads. The analysis successfully identified and quantified 12,688 phosphorylation sites, of which 11,398 were localized with high probability (site probability>0.85) and identified in 3≥ samples for at least one condition. Among these were 2.44% (278) tyrosine, 19.53% (2226) threonine, and 78.03% (8894) serine phosphorylation sites, respectively (Figure 2.E and Table S2). The identified phosphorylation sites represented 2,587 proteins in total (Figure 2.E). The strongest correlation (Pearson correlations 0.68-0.94) was found within the biological replicates (Figure S4.A) and median subtraction across biological replicates, resulted in a strengthened correlation based on insulin treatment (Figure S4.B). To gain insights into the phosphorylation site dynamics after stimulation with different insulin concentrations compared to the unstimulated control, differentially regulated phosphorylation sites were analyzed using t-test statistics in volcano plots. These plots depict the fold change in phosphorylation site levels between the insulin-stimulated and unstimulated conditions against the statistical significance (p-value) for each phosphorylation site (Figure S4.C). The 5-minute stimulation with either 0.1, 3, or 100 nM insulin resulted in upregulated levels of 1,297 and 673 phosphorylation sites in insulin-sensitive and -resistant cells, respectively (≥2 fold-change and p<0.05) (Figure 2.F and Table S2).

To characterize the effects of prolonged insulin stimulation with 0.1 nM and 3 nM insulin during the insulin-resistance-inducing protocol on the proteome, two separate analyses were performed to identify changes in proteome abundance associated with insulin sensitivity and resistance. The first analysis served as a “reference proteome” and was based on lysates used to study the IR interactome and phosphoproteome. These cells were lysed with a milder detergent-containing lysis buffer to preserve endogenous protein-protein interactions [57]. Additionally, to improve proteome coverage, a second analysis applied a harsher lysis buffer containing SDS (referred to as “SDS-proteome”) [36], [58]. The reference proteome and SDS-proteome resulted in the identification and quantification of 4,867 and 6,378 proteins, respectively (Figure S5.A, Figure 2.G, and Table S3). The reference proteome displayed good correlation (Pearson correlations between 0.81-0.97) (Figure S5.B) with high similarity between the insulin-sensitive and -resistant cells. The differential expression analysis compared the proteomes from insulin-sensitive and -resistant cells with the assumption of a minimal effect of the 5-minute insulin stimulation on proteome abundance changes. This analysis identified 117 proteins significantly regulated (p<0.05, ≥1.25-fold; downregulation in insulin resistance, 64; upregulation in insulin resistance, 53). (Figure S5.C). The SDS-proteome dataset showed good correlation (Pearson correlation coefficients averaging 0.97) validating the data quality (Figure S5.D). In the differential analysis, 131 proteins were identified with significant abundance changes (FDR<0.05, s0=0.1; downregulation in insulin resistance, 58; upregulation in insulin resistance, 73) (Figure 2.G and Table S3). The study identified proteins differentially regulated between insulin-sensitive and - resistant cells in at least two of three layers of proteomics datasets (Figure 2.H). The limited number of proteins identified as regulated in the proteome, while concurrently exhibiting regulation in the interactome (one in insulin-resistant) or phosphoproteome (6 in insulin-sensitive and 4 in resistant), indicated a low proteome bias in the interactome and phosphoproteome data (Figure 2.H). Noteworthy, proteins regulated in multiple datasets hold importance because they can influence different aspects of cellular signaling, revealing complex layers of cellular regulation. For example, two kinases, EphA2 and serine/threonine-protein kinase B-raf (BRAF), were upregulated in two proteomics layers following the hyperinsulinemia treatment used to induce insulin resistance. Similar observations were made for the proteins: biogenesis of lysosomal organelles complex 1 subunit 3 (BLOC1S3) and Rab11 family-interacting protein 1 (Rab11FIP1), both involved in cellular vesicle trafficking, in insulin-sensitive cells (Figure 2.H). In summary, the study applied MS technology to create three IR-centric datasets, encompassing IR interactors, phosphorylation sites, and protein levels. These datasets form the foundation for a thorough investigation of dose-response insulin signaling in insulin resistance.

### Interactome analysis reveals distinct IR networks in insulin-sensitive and -resistant cells

Considering the significance of protein-protein interactions in modulating cellular signaling, the aim was to gain an understanding of the IR interactome in induced insulin-resistant HepG2 IGF1R KO cells. The approach involved two strategies: first, examining the strongest IR adaptor candidates with >5 annotated peptide precursors and ≥10-fold enrichment compared to the reference proteome (Figure 2.C and Table S1). Second, exploring the dynamic interactome through differentially regulated IR interactors between unstimulated and insulin-stimulated conditions at all three doses for insulin-sensitive and insulin-resistant cells (Figure S6.A-B). To uncover important signaling components in the IR pathway, the top 207 IR interactors were analyzed. These included 25 kinases, 3 SH2 domain proteins, 6 SH3 domain proteins, 11 PH domain proteins, and 3 Cbl-PTB domain proteins (Figure 3.A and Table S1). Enrichment analysis of Kyoto Encyclopedia of Genes and Genomes (KEGG)-terms confirmed the dominance of the insulin signaling pathway, supported by the identification of key interactors such as PIK3R1, AKT, GAB1, and IRS1 (Figure 3.A-B). Combined, these findings validated the IR co-IP data from the endogenous IR-expressing cell model. To identify notable differences in the basal IR interactome between insulin-resistant and -sensitive cells, the interactor peptide precursor ratios were compared for the two cell states. We identified previously unreported IR interactors and observed variations in the interactor ratios between insulin-sensitive and -resistant interactome for several proteins (Figure 3.A). Further experimental validation of these is needed.

**Figure 3.**
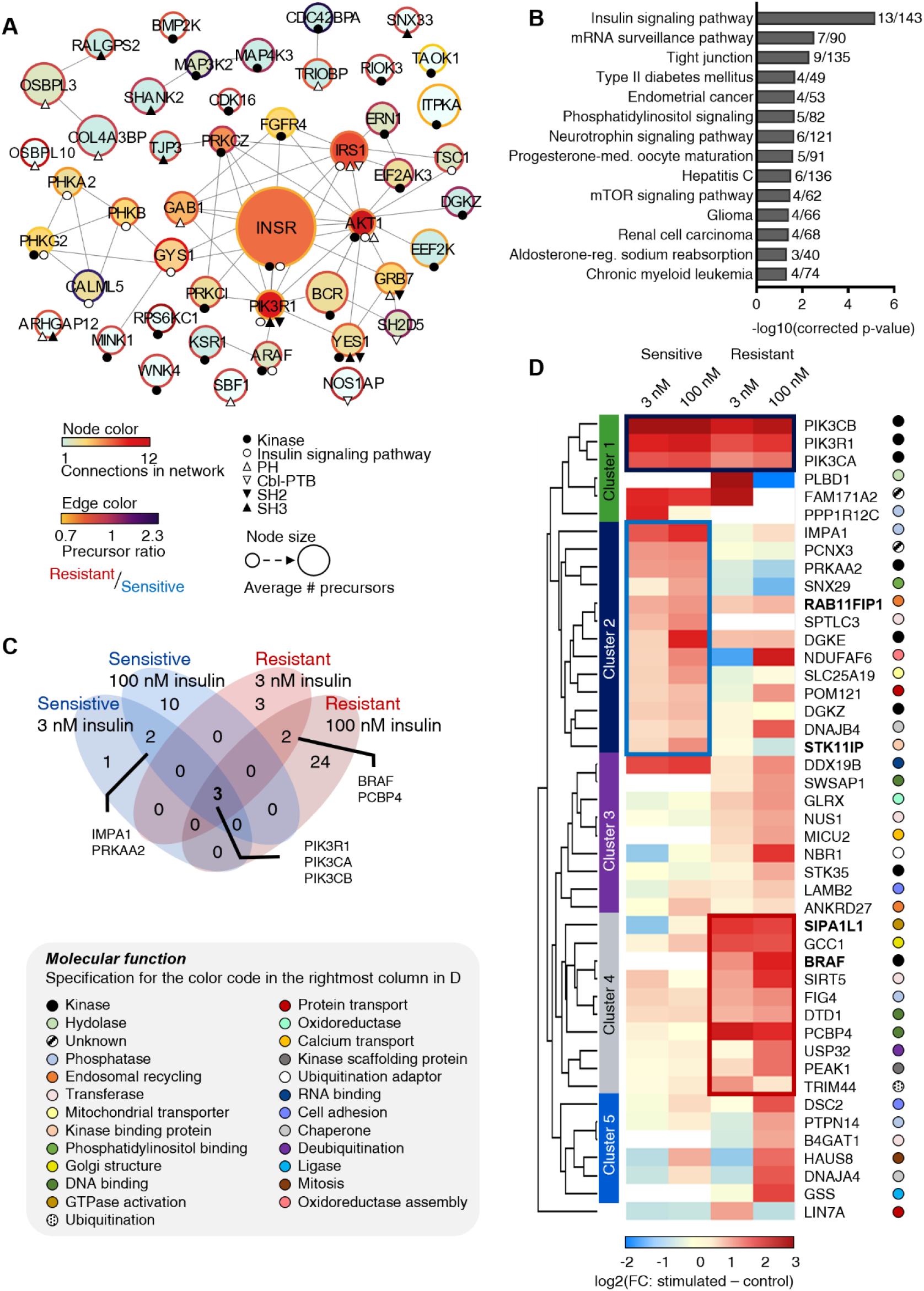
Insulin receptor interactome analysis reveals distinct signaling responses in insulin-sensitive and insulin-resistant cells. **A**. Protein network of a subset of the strongest candidate IR interactors, showing kinases, proteins associated with insulin signaling, and proteins containing specific protein-protein interaction domains (SH2, SH3, PH, and Cbl-PTM domains). Node size indicates the number of annotated precursors, node color reflects connectivity within the network, and edge color represents the ratio of annotated precursors in unstimulated insulin-resistant versus sensitive conditions. **B**. Significantly enriched KEGG pathways for candidate IR interactors (n=207). **C**. Venn diagram showing the overlap of insulin-dependent IR interactors after stimulation with 3 or 100 nM insulin in insulin-sensitive and -resistant cells. Gene names are shown for proteins identified in more than one condition. **D**. Heatmap displaying the fold-change enrichment of the filtered insulin-dependent IR interactome upon stimulation with 3 or 100 nM insulin compared to unstimulated controls in insulin-sensitive and insulin-resistant HepG2 IGF1R KO cells (two-sided t-test, FDR<0.05, s0=0.1) (Figure S6). Interactors significantly recruited in at least one of the conditions are included. Clustered based on hierarchical clustering, divided into 6 clusters (LIN7A in the bottom clustered alone). Cluster 2 (blue frame) and 4 (red frame) display a tendency for stronger interactions in insulin-sensitive and resistant cells, respectively. Protein names in bold denote proteins additionally regulated in the phosphoproteomics dataset (Figure 2.H). Colored circles, right to gene names, denote the primary molecular function of the interactors, with a specification box to the left.

The regulation of dynamic IR interactors was characterized by significant fold-change differences between the 5-minute insulin stimulation and the unstimulated control, as indicated by the respective volcano plots (Figure S6.A-B and Table S1). A separate analysis of the dynamic interactome in insulin-sensitive and -resistant cells was conducted to avoid the potential bias introduced by the decreased IR protein levels in the insulin-resistant state. This approach was taken due to the variability in IR protein levels between the two cell types. The analysis revealed that the PI3K complex, comprising catalytic subunits A and B and a regulatory subunit, showed dynamic recruitment in both insulin-sensitive and -resistant cells with 3 and 100 nM insulin stimulation, but not with 0.1 nM insulin stimulation (Figure 3.C and Figure S6.C). This aligns with the undetected IR and AKT phosphorylation at this insulin concentration (Figure 1.A), indicating minimal activation and signal transduction.

The hierarchical clustering analysis of the significant IR interactors divided the data into six clusters, each displaying distinct behavior in dynamic IR interactors between insulin-sensitive and -resistant cells. Cluster 1 revealed a consistently high level of IR-recruited PI3K complex after 5 minutes of stimulation with 3 and 100 nM insulin in the insulin-sensitive cells, while -resistant cells required 100 nM insulin levels for similar recruitment levels (Figure 3.D). The reduced PI3K recruitment confirmed diminished insulin responsiveness in insulin-resistant cells. Clusters 2-6 showed varying dynamic recruitment of IR interactors in insulin-sensitive and resistant cells based on 5 minutes of 3 or 100 nM insulin stimulation. Cluster 2 contained 13 interactors with a tendency for stronger insulin-induced recruitment in insulin-sensitive compared to resistant cells. The insulin-sensitive state interactors included the two kinases: acetyl-CoA carboxylase kinase (PRKAA2) and diacylglycerol kinase epsilon (DGKE) (Figure 3.D). Also, Rab11FIP1, known for its involvement in receptor tyrosine kinase recycling [59], was detected among these potentially stronger insulin-sensitive interactors, (Figure 3.D). Additionally, phosphorylation levels of Rab11FIP1 were found to be downregulated in insulin-resistant cells compared to insulin-sensitive cells (Figure 1.H and Table S2). In cluster 4, 10 proteins showed a tendency to be more strongly recruited to the IR upon 5 minutes of 3 and 100 nM insulin stimulation in insulin-resistant compared to sensitive cells. This cluster contained the phosphatase polyphosphoinositide phosphatase (FIG4) and, interestingly, the kinase BRAF, which is usually linked with the MAPK signaling pathway. Additionally, BRAF demonstrated higher phosphorylation levels in insulin-resistant cells compared to sensitive cells. While BRAF has not previously been linked to insulin resistance, its kinase activity is regulated by metformin treatment, a first-line therapy to enhance insulin sensitivity, making it an interesting target for further studies [60], [61]. Cluster 5 contained 6 potential interactors showing increased recruitment for 100 nM insulin stimulation in insulin-sensitive and resistant cells, compared to their respective 3 nM stimulation and included the tyrosine-protein phosphatase non-receptor type 14 (PTPN14). The hierarchical clustering analysis showed differences in the recruitment of dynamic IR interactors between insulin-sensitive and -resistant cells. The study found that some dynamic IR interactors had differentially regulated phosphorylation levels, which could be potential targets for further investigation into altered insulin resistance-associated signaling.

### Phosphoproteome analysis uncovers sustained ERK signaling and impaired insulin-stimulated AKT signaling response in insulin-resistant cells

The insulin-resistant phenotype is known to display a reduced activation of the metabolic insulin-induced cellular response [2], [4], [62], [63], which we initially confirmed by showing decreased phosphorylation levels of IR and phosphorylation of AKT (Figure 1.A-C and Figure S1.B). To further characterize the phosphoproteome, the level of regulation of insulin-stimulated phosphorylation sites in insulin-sensitive and -resistant cells was compared. Based on significance testing across all conditions, 1,337 phosphorylation sites showed significantly regulated levels and based on the hierarchical clustering analysis, 10 clusters were defined, resulting in the formation of the large cluster A (Figure 4.A). However, this cluster contained visible sub-clusters that were diverging in their phosphorylation pattern. Cluster A.1 showed decreased phosphorylation levels in unstimulated insulin-sensitive, but not in insulin-resistant cells, compared to their respective insulin-stimulated conditions. The maximum phosphorylation site abundance was reached with the low dose of insulin (0.1 nM) stimulation in insulin-sensitive cells and across all conditions in insulin-resistant cells. In contrast, cluster A.2 displayed an insulin dose-response upregulation in phosphorylation levels for insulin-sensitive and -resistant cells, with a faster increase and higher maximum intensity observed in insulin-sensitive cells (Figure 4.A). A kinase motif enrichment analysis of the amino acid sequence preferences adjacent to phosphorylation sites in cluster A.1 showed an overrepresentation of proline (P) at the +1 position, which is the consensus motif for proline-directed kinases like ERK1/2 kinase substrates [64]. The prolonged activation of ERK/MAPK, resulting from the elevated insulin concentration in the insulin-resistance-inducing protocol, persisted in the control condition without being attenuated, causing phosphorylation of ERK1/2 kinase substrates in the insulin-resistant cells (Figure 4.B). From a similar analysis of cluster A.2, peptides with arginine (R) at position -3 relative to the phosphorylated site, which is the consensus motif for AKT kinase substrates [64], were enriched. Additionally, the recognition sequence for basophilic protein kinases like AKT, p70, S6 kinase, and RSK includes R residues at both positions -3 and -5 [64], partially evident in the cluster motif (Figure 4.B). This observation confirmed the reduced AKT response in insulin-resistant cells, and it was intriguing that the analysis showed differential AKT and ERK substrate motif phosphorylation upon insulin stimulation in insulin-sensitive and -resistant cells, in combination, defining the insulin-resistant state. In addition to the cluster analysis, the regulated phosphorylation sites in insulin sensitivity and resistance were globally compared, focusing on high-concentration stimulations (Figure 4.C). KEGG pathway enrichment analysis further supported the upregulation of terms associated with endocytosis and MAPK pathway signaling in the insulin-resistant state (Figure 4.D). These findings are intriguing, given the observed differences in IR levels and ERK1/2 kinase substrate phosphorylation in insulin-sensitive and - resistant cells.

**Figure 4.**
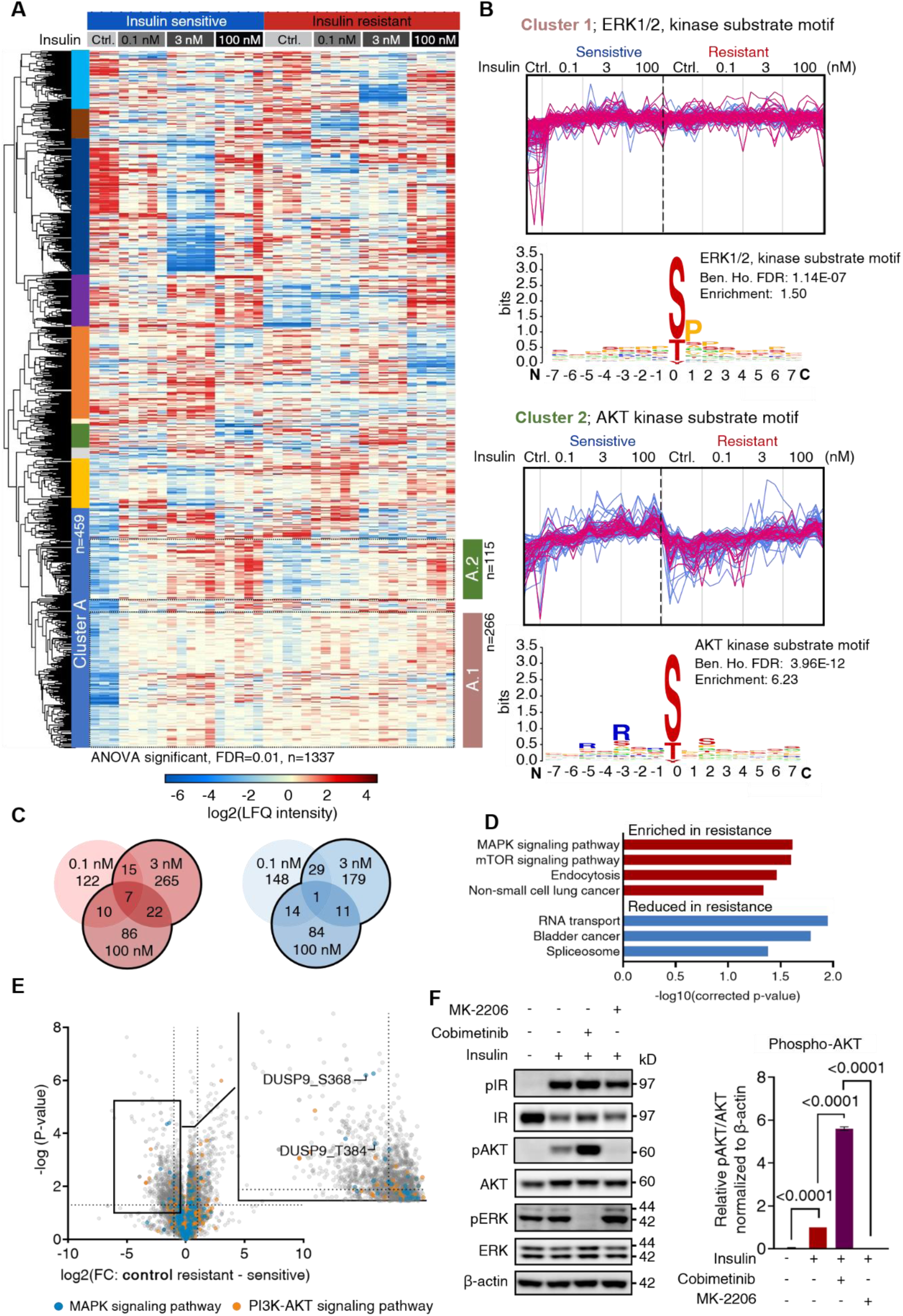
Phosphoproteomics reveals sustained ERK signaling and regulation of DUSP9 phosphorylation in insulin resistance. **A**. Hierarchical clustering of significantly regulated phospho-site levels in response to stimulation with 0.1, 3, or 100 nM insulin in insulin-sensitive and insulin-resistant cells (n=1337; ANOVA; FDR<0.01, s0=0.1). With 10 clusters defined from hierarchical clustering, shown as left color-bar, and subclusters of A, shown to the right. **B**. Cluster profile and enriched sequence motif analysis for subcluster A.1 and A.2 shown in A, for motifs with strongest enrichment and lowest Benjamin-Hochberg (Ben. Ho.) FDR-corrected p-values. The profiles for phosphorylation sites with ERK (top) and AKT (bottom) sequence motifs are highlighted in purple. **C**. Venn diagram showing numbers of phosphorylation sites with induced (red) or decreased (blue) levels in resistant cells upon stimulation with 0.1, 3, or 100 nM insulin compared to unstimulated cells (2-fold change, p<0.05). **D**. KEGG pathway enrichment analysis of upregulated phosphoproteins following 3 or 100 nM insulin stimulation in insulin-sensitive and -resistant cells (2-fold change, p<0.05). **E**. Volcano plot showing -log10(p-value) against log2(fold change) of phosphorylation site LFQ intensity for unstimulated insulin-sensitive and -resistant cells. Phosphorylation sites from proteins involved in the MAPK and PI3K-AKT signaling pathways are highlighted in blue or orange, respectively. Zoomed view highlights phosphorylation sites downregulated in insulin resistance from the MAP kinase phosphatase, DUSP9 (dotted line: ≥2-fold change and significance, p<0.05). **F**. Immunoblot and quantification of HepG2 IGF1R KO cells stimulated for 24 hours with 3 nM insulin without or with inhibition of AKT (MK-2206) or MEK (cobimetinib). Representative blot of n=3 independent biological replicates with p-values<0.05 annotated (two-sample unpaired t-test).

Given the non-attenuated ERK1/2 kinase substrate phosphorylation in insulin resistance, in unstimulated conditions, the differential regulation of phosphorylation sites was examined at the basal level in the two conditions (Figure 4.E). The level of regulation between unstimulated insulin-sensitive versus -resistant cells was evaluated by differential phosphorylation site analysis. From the resulting volcano plot, reduced phosphorylation levels at two sites in dual-specificity protein phosphatase 9 (DUSP9) (Ser368 and Thr384) were identified in insulin-resistant cells (Figure 4.E). Noteworthy, only insulin-resistant cells showed upregulated phosphorylation levels of DUSP9 Ser368 and Thr384 upon insulin stimulation (>1.5-fold at 3 nM and >2-fold at 100 nM, p<0.05) (Figure S4.C). DUSP9 dephosphorylates MAPKs, including ERK1/2 [65], and several studies have shown its involvement in insulin resistance [66]–[68]. Thus, it was speculated whether the lowered phosphorylation levels of DUSP9 at Ser368 and Thr384 under unstimulated insulin-resistant conditions, potentially impact the activity or stability of the protein, leading to sustained ERK phosphorylation and signaling. Additionally, considering the differential ERK and AKT signaling responses observed in insulin-sensitive and -resistant cells, the signaling response was analyzed by inhibiting these two pathways. HepG2 IGF1R KO cells were treated with 3 nM insulin alone or after pre-treatment with either cobimetinib (an inhibitor of mitogen-activated extracellular kinase (MEK)) [69] or MK-2206 (an AKT inhibitor) [70], [71]. Immunoblot analysis confirmed the ability of cobimetinib and MK-2206 to reduce the activation of ERK and AKT, respectively. Intriguingly, inhibiting the MAPK signaling pathway led to a ≥5-fold increase in AKT phosphorylation, strongly suggesting potential cross-talk between AKT and ERK in insulin signaling. This finding further emphasizes the potential significance of the observed differential DUSP9 phosphorylation in our insulin-resistant model (Figure 4.F).

### Proteome analysis reveals upregulation of EphA2 expression in insulin-resistant cells and identifies this gene as insulin-regulated

To gain a deeper understanding of the insulin-resistant HepG2 IGF1R KO cell model in terms of proteome remodeling, we performed volcano plot analysis of the SDS- and reference-proteomes to identify significant fold-changes in protein levels between insulin-sensitive and - resistant cells (Figure 5.A and Figure S5.C). An enrichment analysis of KEGG-terms was performed to characterize the proteins with abundance changes. The analysis showed that proteins involved in MAPK signaling and axon guidance pathways were upregulated, while proteins related to glycolysis, gluconeogenesis, glucose, and amino acid metabolic pathways were downregulated in insulin resistance (Figure 5.B).

**Figure 5.**
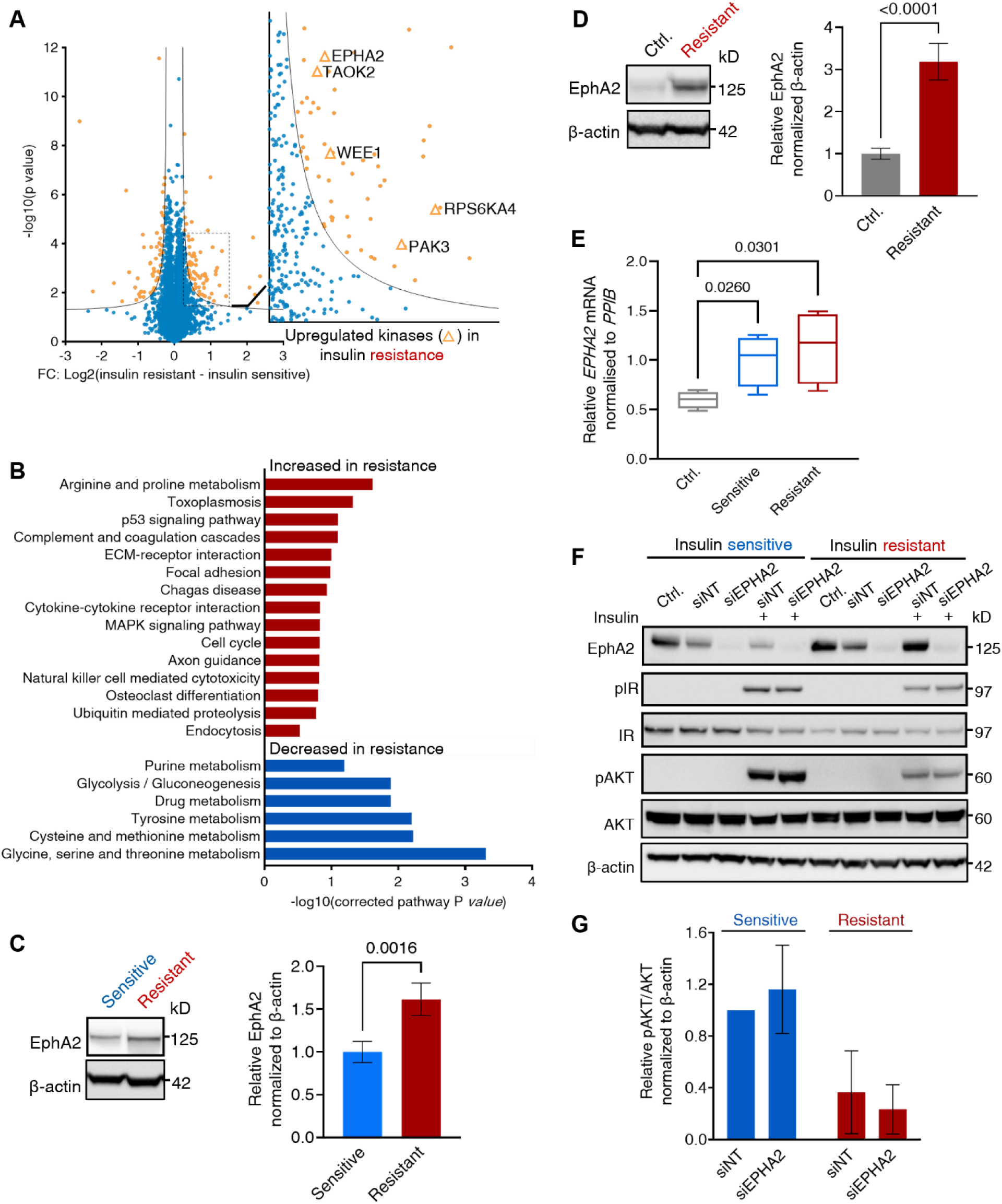
Proteome analysis shows upregulated EphA2 expression levels in insulin-resistant cells and uncovers it as an insulin-inducible gene. **A**. Volcano plot presenting differentially regulated proteins in insulin-sensitive and resistant SDS-based cell lysates from single-shot proteome MS analysis. Significantly regulated proteins are highlighted in orange (FDR<0.05, S0 = 0.1). Zoomed view shows upregulated proteins in insulin-resistant cells, with kinases labeled with gene names (n=4 independent biological experiments). **B**. KEGG term analysis of proteins significantly upregulated (red) or downregulated (blue) in insulin resistance. **C**. Immunoblot and quantification of EphA2 abundance in lysates from insulin-sensitive and -resistant cells (n=4 independent biological experiments). **D**. Immunoblot and quantification of EphA2 in unstimulated cells (48-hour serum-starved control; ctrl.) or cells subjected to the insulin resistance-induction protocol (including 48-hour 3 nM insulin treatment). **E**. qPCR results presenting the mRNA levels of EPHA2 in HepG2 IGF1R KO cells treated as specified in D. Median of six technical replicates for n=4 biological independent replicates. **F**. Representative immunoblot of insulin-sensitive and resistant HepG2 IGF1R KO cell lysates with EphA2 siRNA knock-down (KD) (NT=non-targeting control). Cells stimulated with 3 nM insulin are indicated (n=4 biological independent replicates) (two-sample unpaired t-test). **G**. Quantification of phospho-AKT normalized to AKT and β-actin levels from immunoblot, shown in F, in insulin-sensitive and -resistant lysates with and without EphA2 KD. Relative to siNT in the insulin-sensitive cells.

In the SDS-proteome, five protein kinases were identified among 6,378 proteins with upregulated protein levels in insulin-resistant compared to -sensitive cells. They were: EphA2, serine/threonine-protein kinase TAO2 (TAOK2), wee1-like protein kinase (WEE1), ribosomal protein S6 kinase alpha (RPS6KA), and serine/threonine-protein kinase PAK 3 (PAK3) (Figure 5.A and Table S3). Of these kinases, the reference-proteome only confirmed EphA2 to be upregulated in insulin-resistant cells, and thus, a shared significant finding in the two proteome analyses (Figure S5.C and E). The RTK EphA2 belongs to the Eph receptor family and is known to play a role in mediating cell-cell communication and regulating various cellular processes [25]– [27]. However, to our knowledge no information exists on its role in insulin resistance or gene expression regulation by insulin. By immunoblotting, the ∼1.5-fold increase in EphA2 abundance at the protein level in insulin-resistant compared to insulin-sensitive cells was confirmed (Figure 5.C). Importantly, EphA2 levels were increased upon insulin stimulation at protein and mRNA levels independently of the insulin resistance, and this underscored the role of insulin signaling-dependency of these changes (Figure 5.D-E). Moreover, increased phosphorylation levels were identified of EphA2 Tyr594 in the 3 nM insulin-stimulated insulin-resistant cells compared to the - sensitive in the phosphoproteome (Figure 2.H, Figure S7.A). The insulin-stimulated phosphorylation was investigated in insulin-sensitive and -resistant cells, separately. The level of an EphA2 doubly phosphorylated peptide covering the autophosphorylation sites Tyr588 and Tyr594 was downregulated in response to 0.1 nM insulin stimulation in insulin-sensitive, and for all three insulin concentrations in the -resistant cells (Figure S4.C and Table S2).

Based on these results, showing differential levels of EphA2 in insulin-sensitive and insulin-resistant cells, it was speculated that increased EphA2 levels could be causally involved in the insulin-resistant phenotype by suppressing the ability of activated IR to phosphorylate AKT. Therefore, to elucidate its potential role in insulin resistance, a small interfering RNA (siRNA) knockdown (KD) experiment was performed targeting EphA2 in both cell states. The immunoblot showed effective KD of EphA2 in the HepG2 IGF1R KO cell model, however, it did not restore insulin-stimulated phosphorylation levels of IR and AKT in the insulin-resistant cell model (Figure 5.F-G). These results confirmed a strong influence of insulin stimulation on increased EphA2 levels, and aligned well with insulin resistance, as an outcome of prolonged cell culturing with high insulin concentration according to the resistance-inducing protocol. However, the regulation of EphA2 was independent of insulin resistance. To our knowledge, this is the first reporting of EphA2 as an insulin-inducible gene.

### Expression level of IGF1R correlates with regulation of EphA2 by insulin stimulation in different cell lines

Since EphA2 KD could not restore AKT signaling in insulin-resistant cells to the level in - sensitive cells, we assumed that the upregulation of EphA2 did not play a major causal role in resistance. However, we speculated whether the insulin-dependency of EphA2 expression was a general finding across other cell types or specific to hepatocyte-like cells. Therefore, HepG2 WT cells were treated with 3 and 100 nM insulin for 24 hours (Figure 6.A-B). Interestingly, unlike in HepG2 IGF1R KO cells, insulin stimulation did not induce expression of EphA2 in HepG2 WT cells, indicating an inhibitory or rewiring role of IGF1R in this response. Therefore, a total of 11 insulin-responsive cell lines were screened: 4 hepatoma (HepG2 WT, HepG2 IGF1R KO, Hep3B, and H4IIE), 3 acute myeloid leukemia (AML) (MOLM-13, THP1, and EOL-1), and 4 breast cancer (BT549, BT474, HCC38, and HCC197) cell lines for insulin-induced EphA2 regulation. The cells were chosen based on their differential IR-to-IGF1R expression levels (Figure 6.D). The IR-to-IGF1R ratio is generally higher in hepatic and AML cells, while breast cancer cells show varying but lower ratios given higher IGF1R expression levels (Figure 6.C-D). For the hepatoma cell lines, insulin stimulation induced EphA2 expression in HepG2 IGF1R KO and the rat hepatoma H4IIE cells known to lack IGFIR expression [72]. H4IIE cells showed dose-response induction, but not in HepG2 WT and Hep3B cell lines. Among the AML cell lines, ∼5-fold increase of EphA2 protein level with 3 nM insulin stimulation was observed for MOLM-13, whereas THP1 or EOL-1 cells showed no EphA2 induction. None of the screened breast cancer cell lines showed EphA2 protein regulation after insulin stimulation (Figure 6.A-B). Taken together, HepG2 IGF1R KO, H4IIE, and MOLM-13 cells exhibited the lowest IGF1R protein levels, supporting the hypothesis of a low IGF1R expression requirement for insulin-induced EphA2 upregulation (Figure 6.C-D). Previous studies have demonstrated that high insulin concentrations can activate IGF1R, initiating proliferative signaling activities [73]–[75]. Additionally, heterodimers between IR and IGF1R can form, and these hybrid receptors bind insulin with reduced affinity compared to IR and have been linked to diabetic individuals [76], [77]. This underscores the importance of understanding the interplay between insulin signaling via these two homologous growth factor receptors, IGF1R and IR, in both metabolic and growth-related contexts.

**Figure 6.**
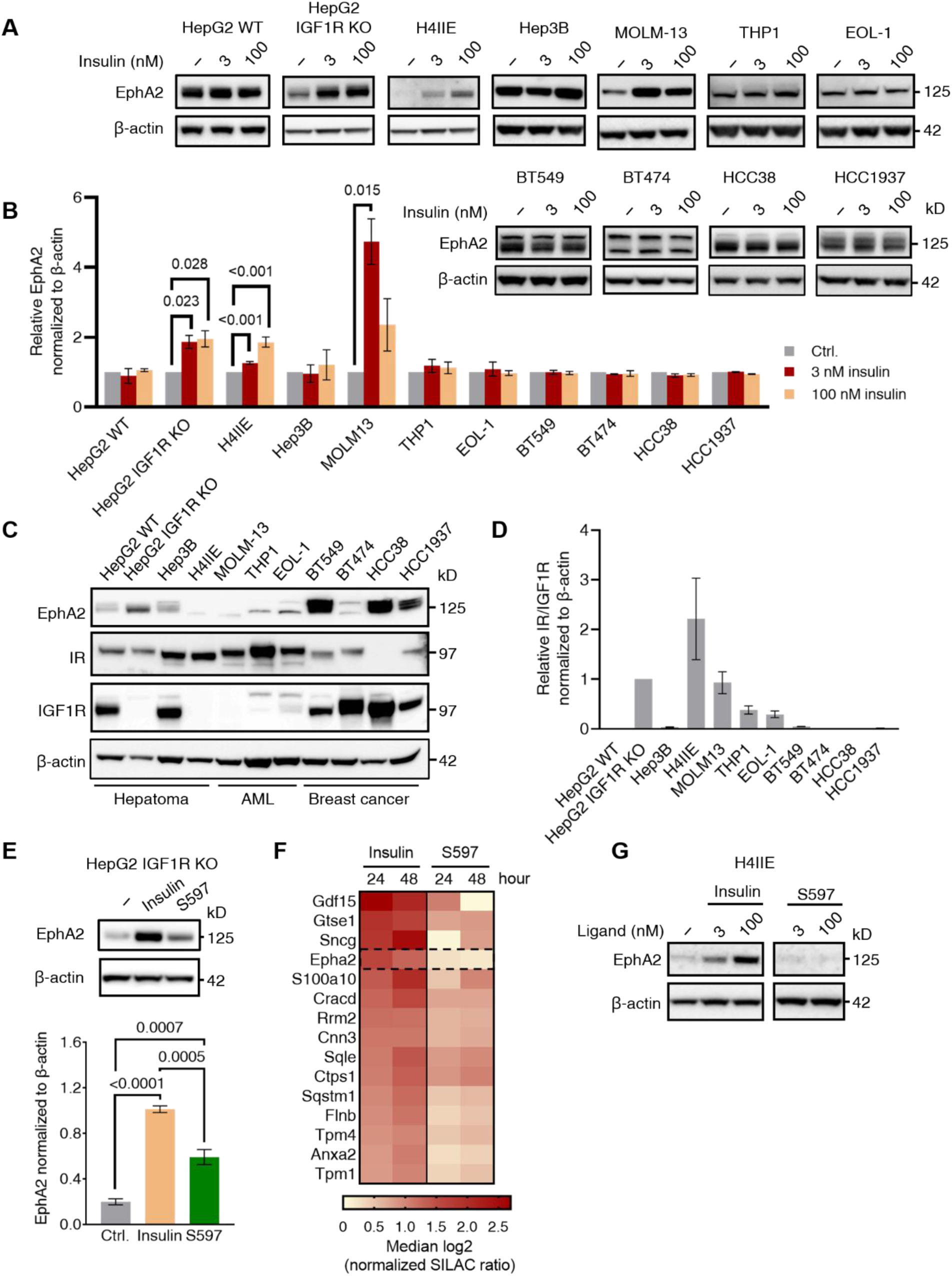
Induction of EphA2 by insulin correlates with a high IR-to-IGF1R ratio and the insulin agonist 597 does not show the same EphA2-inducing characteristics. **A**. Immunoblot of EphA2 in a panel of screened hepatoma, breast cancer, and acute myeloid leukemia (AML) cell lines serum-starved or treated with 3 or 100 nM insulin for 24 hours. Representative of n=2 (n=3 for H4IIE and Hep3B) biological independent experiments. **B**. Quantification of immunoblot in A. EphA2 protein levels after insulin stimulation relative to untreated (ctrl.) for individual cell lines (two-sample unpaired t-test). Normalized to β-actin levels. **C**. Immunoblot of EphA2, IR, and IGF1R in the cell panel shown in A and B. Representative of n=2 biological independent replicates. **D**. Quantification of the relative IR-to-IGF1R ratios from immunoblot in C, normalized to β-actin levels. **E**. Immunoblot and quantification of EphA2 in HepG2 IGF1R KO cells untreated or treated with 100 nM insulin or the partial IR agonist S597 for 24-hours (two-sample unpaired t-test) (n=4 independent biological experiments). **F**. Heatmap of median log2 protein SILAC ratios for significantly insulin-induced protein levels in H4IIE cells (significance B testing, FDR<0.05). Cells were cultured with 100 nM insulin or S597 for 24 or 48 hours (n=3 independent biological experiments). **G**. Representative immunoblot of EphA2 in H4IIE cells treated for 24 hours with 100 nM insulin or S597 (two-sample unpaired t-test) (n=3 independent biological replicates).

Based on the phosphoproteome data showing a sustained level of ERK signaling in the insulin-resistant cells, this led to the question of whether the balance between PI3K-AKT and MAPK downstream signaling impacted the regulation of EphA2. Therefore, HepG2 IGF1R KO cells were stimulated for 24 hours with 3 nM and 100 nM or the partial IR agonist S597, which has been shown to primarily signal via the PI3K-AKT pathway [29]. Intriguingly, stimulation with S597 did not result in upregulated EphA2 expression at the protein level, compared to stimulation with insulin. This supported the dependency of induced ERK signaling, in the insulin-induced upregulation of EphA2 (Figure 6.E). To confirm the observed regulation of EphA2 protein levels by insulin but not S597 in another hepatic cell model, the rat hepatoma cell line H4IIE was used. This cell line lacks endogenous IGF1R expression, hence without manipulation mimicking physiological healthy hepatocytes (Figure 6.C-D) [31]. In addition to validating the observations in another cell system, the broader changes in protein abundance following stimulation with insulin and S597 were explored. Therefore, a SILAC experiment in H4IIE cells was performed and cells were stimulated with 100 nM insulin or S597 for 24 and 48 hours followed by an MS-based proteome analysis (Figure S8.A-B). A total of 7,508 proteins were confidently identified and EphA2 protein levels were upregulated after 24 and 48 hours of insulin stimulation, ranking EphA2 as the fourth most induced protein. In contrast, S597-stimulation did not lead to detectable upregulation of EphA2 protein levels, which was verified by immunoblotting (Figure 6.F-G). This supported our hypothesis that insulin regulates EphA2 induction via activation of the MAPK signaling pathway.

### MAPK signaling drives insulin-induced EphA2 expression in cell models with a high IR/IGF1R ratio

To validate that S597 primarily signals through the PI3K-AKT pathway and further elucidate potential differences in downstream signaling by insulin and S597, a SILAC-based phosphoproteome analysis was performed in H4IIE cells. The cells were stimulated with 100 nM insulin or S597 for 5, 15, or 30 minutes before lysis and MS analysis. We identified a total of 11,995 class I phosphorylation sites, each showing a strong correlation across the different conditions (Figure S8.C-D and Table S4). A heatmap of phosphorylation sites with differentially regulated SILAC ratios (≥2-fold in at least one condition), highlighted an enriched cluster associated with the insulin signaling pathway (Figure 7.A). In cluster 1, representing phosphorylation sites within proteins associated with the insulin signaling pathway, phosphorylation linked to the AKT-mediated insulin response exhibited similar patterns for both ligands as shown by Jensen *et al*. [29]. Notably, S597-stimulated cells displayed more pronounced IR Y1186 phosphorylation, confirming IR activation by S597 (Figure 7.B). Furthermore, elevated levels of MAPK phosphorylation (ERK2 Thr183 and Y185; ERK1 Thr203 and Y205) were observed following insulin stimulation, which was less pronounced with S597, aligning with the previous findings on S597-mediated signaling [29] (Figure 7.B).

**Figure 7.**
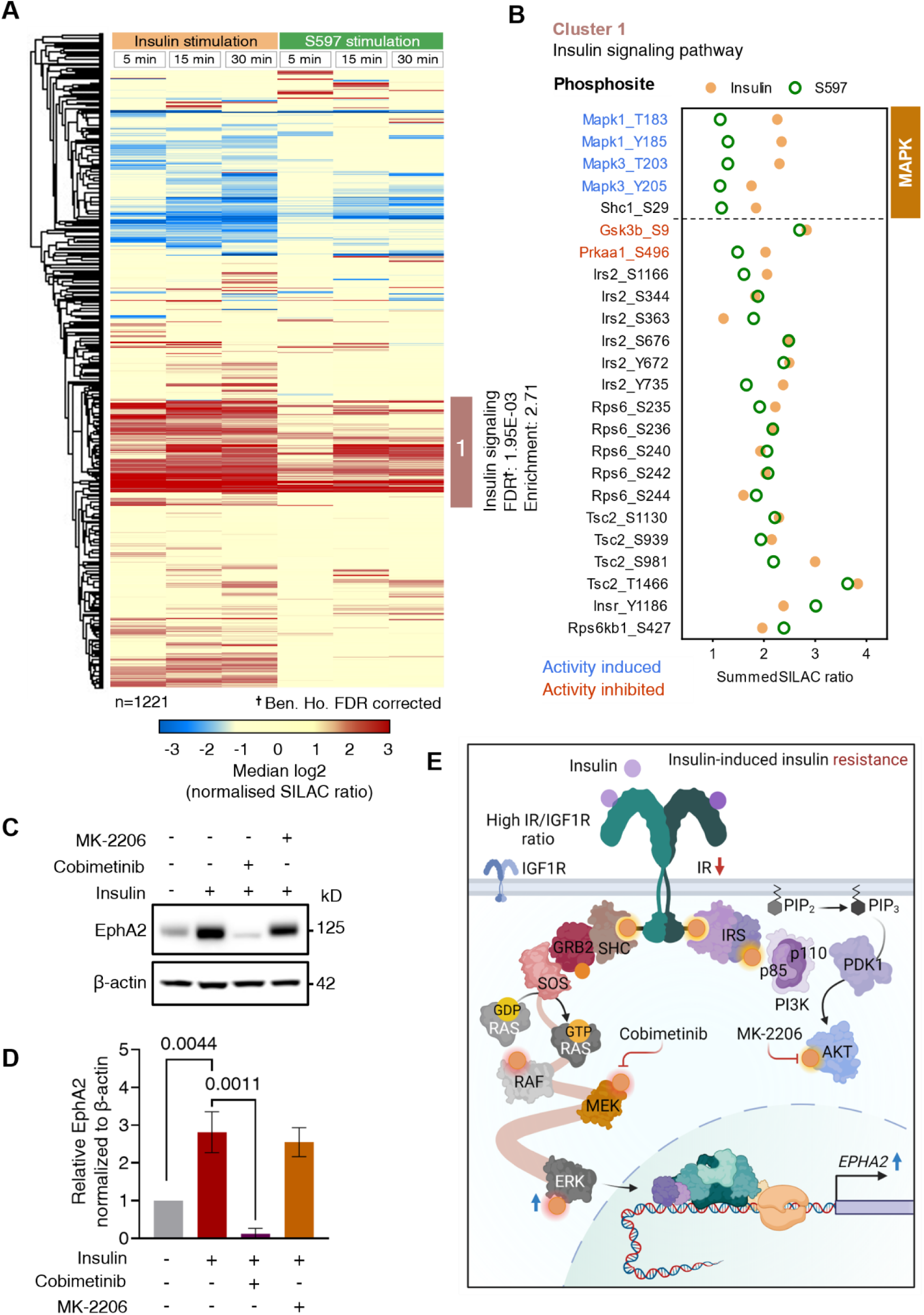
SILAC phosphoproteomics validates primary AKT-directed response by S597 and reveals insulin-mediated EphA2 expression via the MAPK pathway. **A**. Heatmap displaying phosphorylation site SILAC ratios regulated ≥2-fold after 5-, 15-, or 30-minute stimulation with 100 nM insulin or S597 in H4IIE, in at least one condition. The cluster with the overrepresented KEGG insulin signaling pathway is marked. Median SILAC ratios from n=2 independent experiments. **B**. Plot illustrating KEGG insulin signaling pathway phosphorylation sites (labeled cluster in A). Median SILAC ratio, averaged for all time points for insulin (solid yellow) and S597 (hollow green). Regulatory sites are color-coded based on whether phosphorylation induces (blue) or inhibits (red) protein activity. **C**. Immunoblot of HepG2 IGF1R KO lysates treated for 24 hours with 1 µM MK-2206 (AKT) inhibitor, 1 µM cobimetinib (MEK) inhibitor, and 3 nM insulin. Immunoblots of pIR, IR, pAKT, AKT, pERK, and ERK are shown in Figure 4.D. **D**. Quantification of EphA2 protein levels, based on immunoblot in C. Normalized to β-actin levels. **E**. Schematics depicting dysregulation of MAPK signaling in insulin resistance showing the connection to induced EphA2 expression levels and INS/IGF1R ratio dependence.

The reduced ERK signaling and lack of EphA2 induction upon stimulation with S597 support the hypothesis of MAPK-mediated EphA2 regulation by insulin in the IGF1R-deficient cell line models. To further test this hypothesis, EphA2 expression in HepG2 IGF1R KO cells treated with insulin alone or after pre-treatment with either the MEK inhibitor cobimetinib or the AKT inhibitor MK-2206 were examined, using the same lysates employed to evaluate the interaction between AKT and ERK phosphorylation levels in insulin signaling (Figure 4.E). Immunoblot data confirmed a lack of insulin-induced EphA2 expression in cells subjected to MEK inhibition, in contrast to cells treated with insulin alone or in combination with AKT inhibitor (Figure 7.C-D). This confirmed that induction of EphA2 gene expression by insulin is dependent on MAPK pathway activity in this IGF1R-deficient cell model.

## Discussion

In this study, we employed a multi-layered proteomics approach to characterize insulin signaling in insulin-sensitive and resistant conditions in a hepatocellular context. We adapted the insulin resistance-inducing protocol established by Dall’Agnese *et al*. [17] but utilized HepG2 IGF1R KO cells, which as healthy hepatocytes do not express IGF1R [32], in contrast to the HepG2 WT. Although IR and IGF1R share significant homology and engage similar signaling pathways, they show differences in their primary physiological response. The IR primarily regulates metabolic processes, while IGF1R primarily controls growth [73]. The HepG2 IGF1R KO and WT cell models had reduced metabolic AKT signaling during insulin resistance, a defining feature of insulin resistance, but showed differences in other parameters. These differences indicate an interesting interplay between IR and IGF1R, and three examples from our study, where the absence of IGF1R is important for the outcome of insulin signaling, will form the basis for this discussion.

In the insulin-resistant condition, we observed a decrease in overall and surface IR levels, which interestingly differed from the results in HepG2 WT cells [17]. While inconsistent findings exist, studies indicate that chronically elevated insulin levels may result in reduced levels of IR in insulin-resistant and diabetic systems [13]–[15], [54]. Different mechanisms are involved in the reduced IR plasma membrane levels, including clathrin-mediated IR endocytosis and degradation upon internalization [55], [56]. In the IR interactome analysis, we identified the Rab11FIP1 protein, which is implicated in intracellular vesicle trafficking and RTK recycling [19], [59]. However, whether the identification of Rab11FIP1 is linked to the observed higher IR surface levels in the insulin-sensitive cells, due to altered endosomal sorting of the IR in insulin-sensitive and -resistant cells, requires further analysis. Compared to previous interactome studies, we identified 45 dynamic IR interactors, which is deemed a modest number of IR interactors [20], [22]–[24]. Despite the prevalent use of IR overexpression models in most previous studies, our study showed substantial receptor enrichment and successfully identified established and potential previously unknown IR interactors. These findings underscore the sensitivity of the applied workflow enabling studies in a cellular context with endogenous IR expression that more closely resemble physiological conditions [22]–[24].

Another difference between the HepG2 cells with and without IGF1R KO was the reduced ERK activation in the WT model after 5-minute insulin stimulation in insulin resistance [17]. The IGF1R KO cells showed sustained ERK activation, hindering further activation upon 5-minute insulin stimulation. The observation that insulin resistance did not reduce the ERK response is a common phenomenon in insulin resistance [78]. Based on the diverging observations in the HepG2 WT and IGF1R KO cells, it could be interesting to investigate whether the proliferative insulin response depends on a high IR/IGF1R ratio. Since inhibition of the MAPK-ERK signaling pathway in standard cultured HepG2 IGF1R KO cells resulted in an increased PI3K-AKT activation by insulin, this suggests a cross-talk mechanism between ERK and AKT pathway activation following insulin stimulation. Previous studies have addressed cross-talk between the AKT and ERK signaling pathways, both in general and in the context of insulin resistance [11], [79], [80]. A previous study used modeling to predict that activated ERK can inhibit the insulin-stimulated association between IRS1 and PI3K, thus reducing AKT signaling [12]. This observation aligns with our findings. Based on our results, it can be speculated that suppressing the insulin-stimulated ERK response in insulin resistance, might help restore the activation of the metabolic AKT insulin signaling pathway. Interestingly, a study showed restored insulin sensitivity and glucose tolerance in a diabetic mouse model upon treatment with a MEK inhibitor [81]. This supports the relevance of further exploration of IR agonists, like S597, which has been shown to exhibit biased AKT signaling [29], within the field of insulin resistance research. Furthermore, understanding the mechanisms behind sustained MAPK signaling in insulin resistance is important and adds relevance to our observation of the down-regulated phosphorylation levels of the ERK phosphatase, DUSP9. Increased levels of this phosphatase have been shown to protect against insulin resistance [65], [66], [82]. The potential regulatory role of the significantly regulated C-terminal phosphorylation sites of DUSP9 identified in this study is unknown. However, previous research has shown that the C-terminal phosphorylation of the related DUSP2, containing a homologous phosphatase domain to that of DUSP9, has a protein-stabilizing effect [83], [84]. Further research is needed to elucidate the significance of DUSP9 phosphorylation and its connection to ERK signaling in our model.

A key finding in our study was the upregulation of EphA2 protein levels by insulin through the ERK signaling pathway. This aligns with the findings of Macrae *et al*. [85], who demonstrated that EphA2 gene expression is induced via RAF pathway activation in fibroblast and human breast epithelial cells upon hormone activation and that EphA2 protein levels are inhibited with MAPK pathway inhibition in breast cancer cells [85]. Generally, elevated levels of EphA2 are associated with various cancers [26]–[28]. Interestingly, we showed that insulin-stimulated EphA2 induction was specific to cells with a high IR-to-IGF1R ratio. EphA2 is involved in cell adhesion, migration, proliferation, and tissue development. However, the implications of increased EphA2 levels in the HepG2 IGF1R KO cell model with induced insulin resistance remain unknown. Interestingly, another receptor from the Eph receptor family, ephrin type-B receptor 4 (EphB4), has been shown to interact with, and drive IR degradation after insulin stimulation [56]. Based on our co-IP analysis, an interaction between IR and EpHA2 was not observed, and a similar role to that observed for EphB4 thus cannot be confirmed. However, further experiments are needed to confirm this.

The role of high IR-to-IGFR ratio in the insulin-induction of EphA2 is evidence of the apparent role of IGF1R levels concerning insulin signaling. The ability of IR to form hybrid receptors with IGF1R in cells co-expressing the two receptors adds an intriguing dimension to the already complex interplay of the two receptors [76], [86]. Studies have shown that hybrid receptors have reduced insulin affinity compared to the IR [76], [87]. While their function remains uncertain, increased levels of hybrid receptors have been associated with metabolic and mitogenic dysregulated diseases, such as insulin resistance and certain types of cancer [74], [77], [86]. The differences observed in the HepG2 IGF1R KO cell line compared to the HepG2 WT highlight the relevance of this study when investigating insulin resistance in a liver-specific context. However, our model is still a cell line, and therefore, applying the insulin resistance-inducing protocol to more physiologically mimicking cell models, such as liver organoids, would be an evident next step to gain further understanding of insulin resistance.

In conclusion, this multi-layered proteomics study serves as a comprehensive resource, covering the IR signaling dynamics and global proteome changes in an insulin-sensitive and - resistant hepatocellular cell line model with endogenous IR levels and a KO of IGF1R. Using this integrative proteomics approach, we highlighted the role of ERK signaling on the proteome landscape and found that insulin, independently of insulin resistance, induces ERK-dependent EphA2 expression exclusively in cells with a high IR-to-IGF1R ratio.

## Acknowledgement

S.H.J would like to express gratitude to the technicians M. Okkels, R. S. Ingvorsen, and H. Lykkegaard at Novo Nordisk for their introduction and guidance in operating the different cellular assays used in this work. Their input and support have been of immeasurable value. Thanks to S.H. Rahman for contributing to the generation of the HepG2 IGF1R KO cell line.

Additionally, thanks to the members of the Olsen Group at the Proteomics Program, Novo Nordisk Foundation Center for Protein Research (CPR), for their valuable insights and comments. S.H.J was funded by Innovation Fund Denmark under the Ministry of Higher Education and Science (0153-00008) and the Novo Nordisk STAR Fellowship Program. Work at CPR is funded in part by a donation from the Novo Nordisk Foundation (NNF14CC0001). This work has also been supported by the European Research Council through ERC-Synergy grant 810057-HighResCells. All schematic figures were created with <u>BioRender.com</u>

## Supplementary Figures

**Figure S1.**
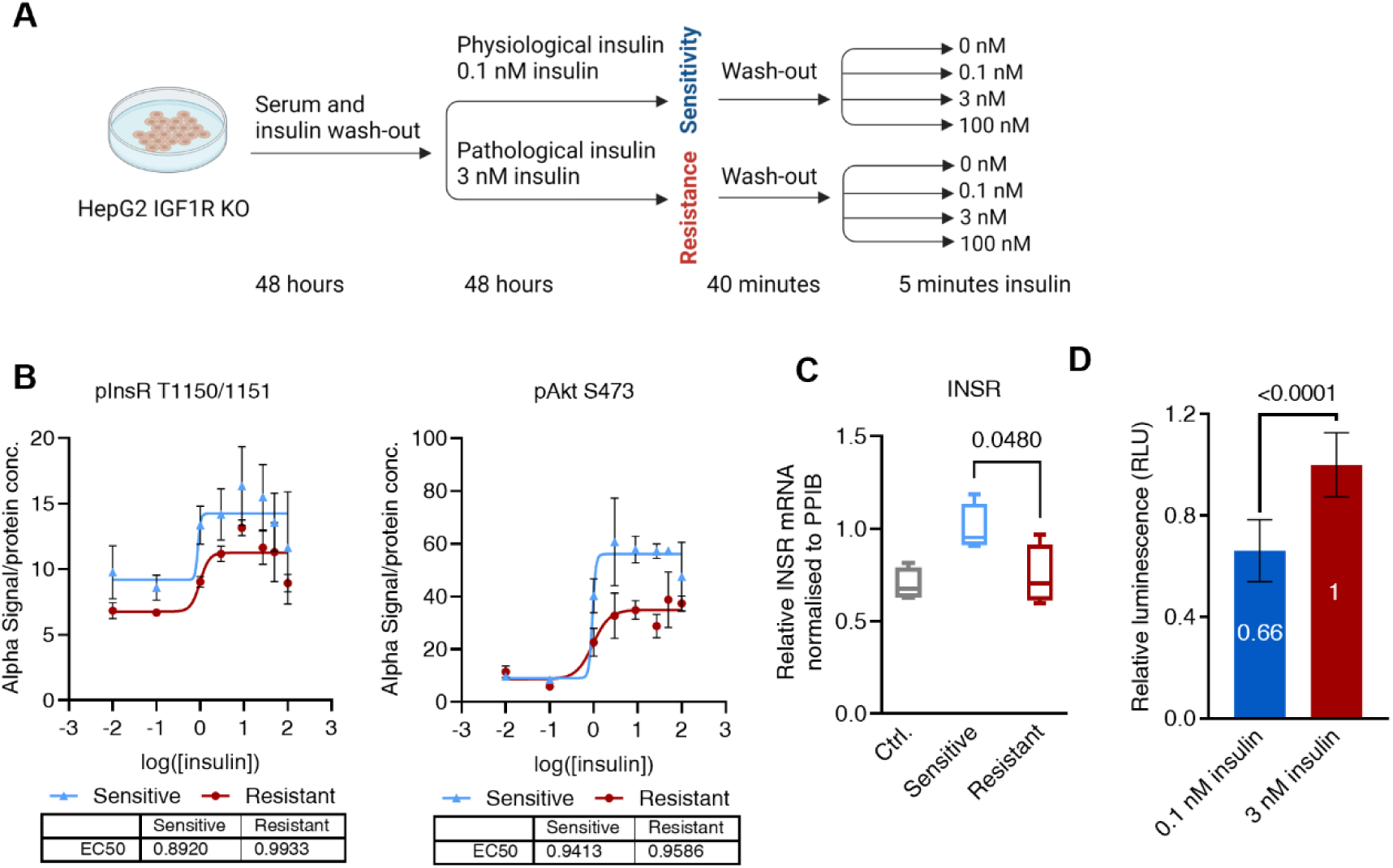
Validation of insulin-induced insulin resistance in HepG2 IGF1R KO cell model. **A**. Schematic of the insulin resistance-inducing protocol in the HepG2 IGF1R KO cell model, adapted from Dall’Agnese et al. [1]. **B**. Insulin dose-response curves of IR and AKT phosphorylation levels for insulin-sensitive and -resistant HepG2 IGF1R KO cells after stimulation with 0-600 nM insulin. The calculated EC50 values are indicated below the curves. Representative example of n=3 independent biological experiments with 3 technical replicates. **C**. qPCR of INSR mRNA levels in HepG2 IGF1R KO cells treated as stated in the insulin-resistance inducing protocol. Control (ctrl.) 48-hour serum starvation. The mean of n=4 biological independent experiments with n=6 technical replicates relative to the insulin-sensitive condition (two-sample unpaired t-test). Only significant p-values are shown. **D**. Relative cell viability was measured in cells cultured for 48 hours with 0.1 nM (physiological) or 3 nM insulin (pathophysiological) after 24-hour serum starvation. Cell viability was determined using the CellTiter-Glo® assay. The relative means (annotated in the respective bars) ±SD of n=3 technical replicates for n=3 biological independent experiments (two-sample unpaired t-test).

**Figure S2.**
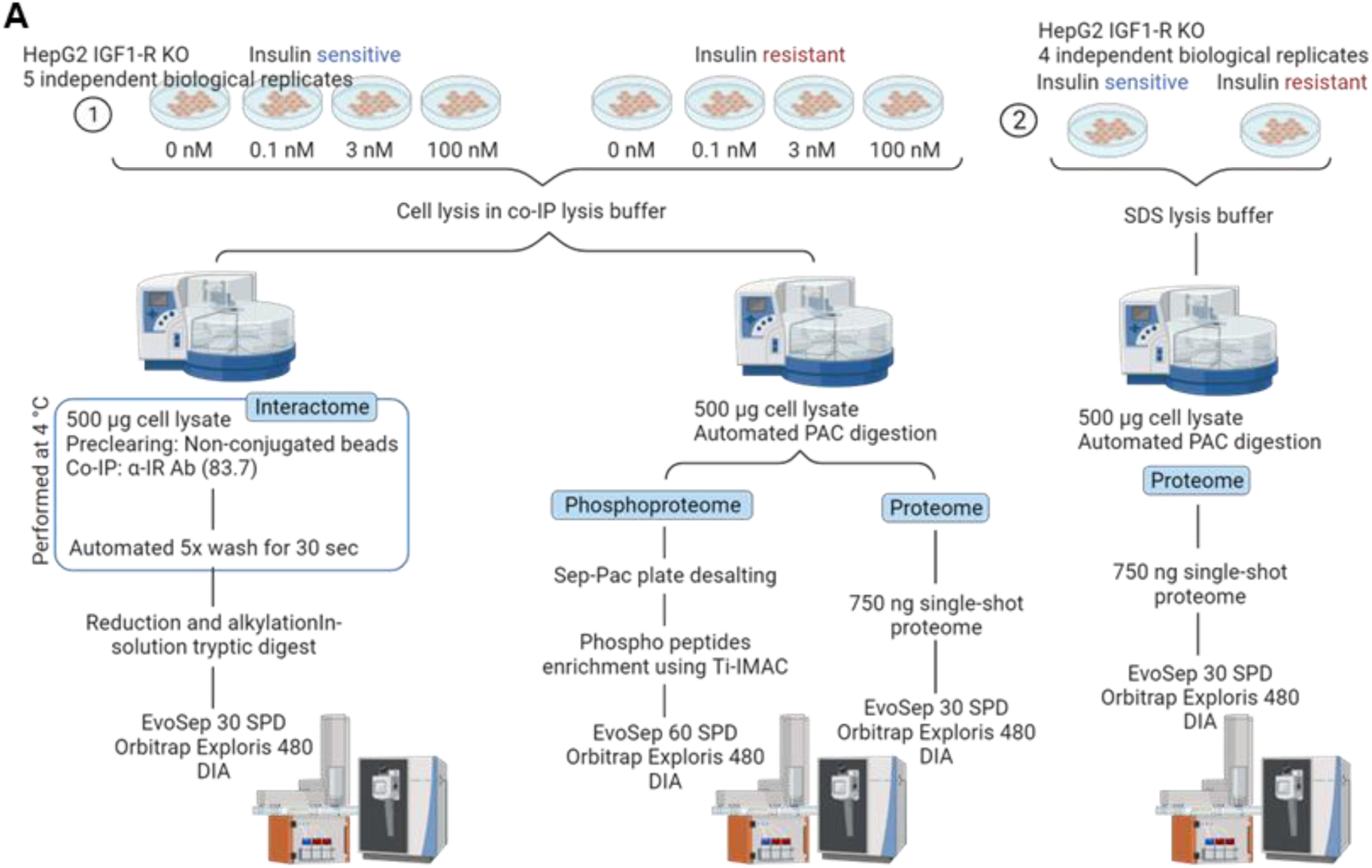
MS-based experimental workflow. **A**. Schematic of the detailed workflow of DIA MS-based analysis for the interactome, phosphoproteome, and reference single-shot proteome in the insulin-resistant HepG2 IGF1R KO cell model (n=5 independent biological replicates). In (2): Workflow for deep-proteome analysis (n=4 independent biological replicates).

**Figure S3.**
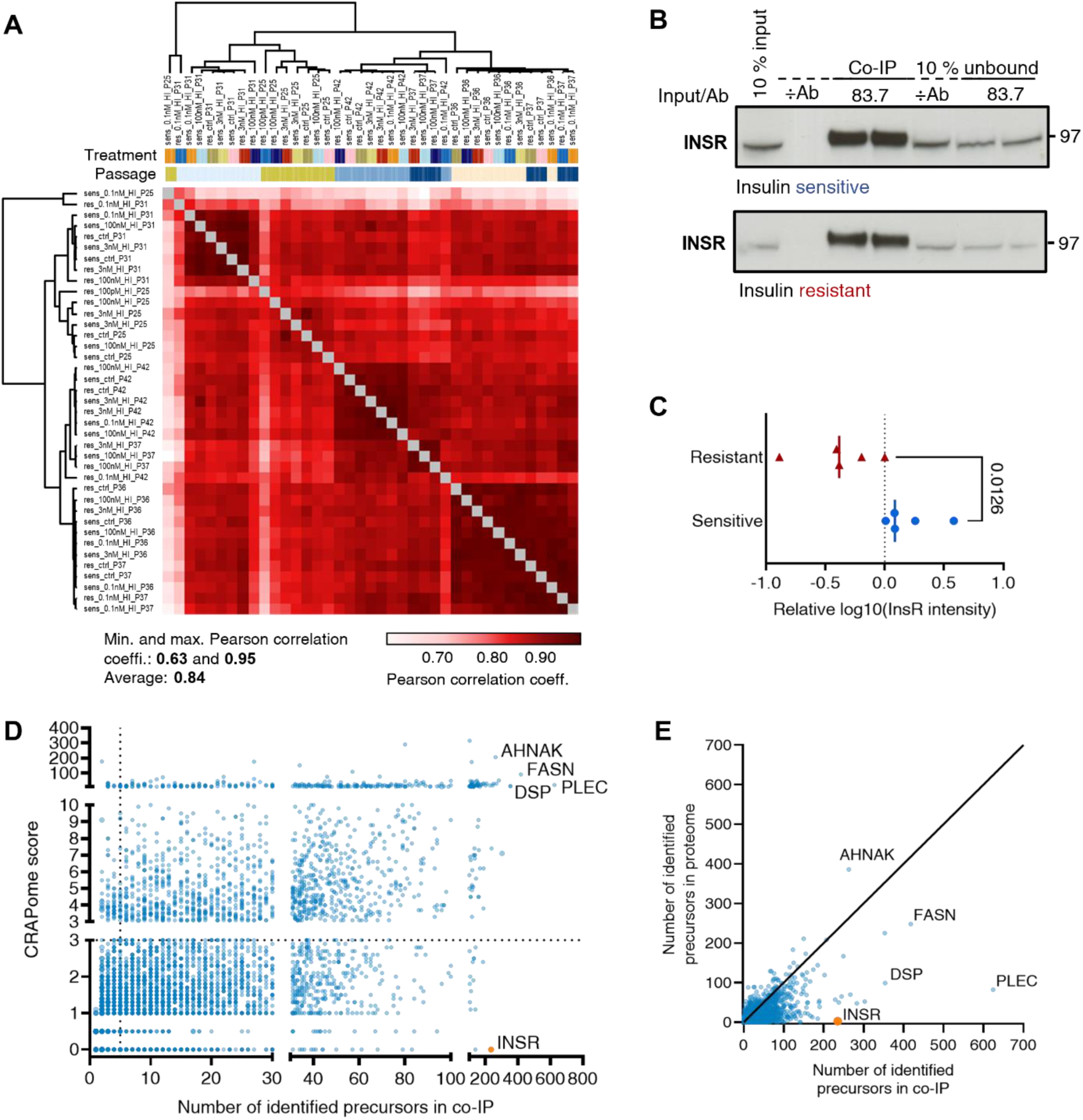
Quality control of interactome data. **A**. Heatmap showing Pearson correlation coefficients of the IR co-IP DIA MS datasets from the insulin-sensitive and -resistant HepG2 IGF1R KO cell model (n=5 biological independent replicates for each condition). The samples are ordered by hierarchical clustering using the Pearson distance metric algorithm. Minimum, maximum, and average Pearson correlation coefficients are indicated. **B**. Immunoblot analysis of IR co-IP in insulin-sensitive and -resistant HepG2 IGF1R KO cell lysates. Control without antibody (÷Ab) for co-IP was included. **C**. -log10-transformed MS-intensities of IR from the co-IP MS dataset (after median subtraction on cell passage) in insulin-sensitive and -resistant cells without insulin stimulation (two-sample unpaired t-test). **D**. Scatter plot depicting the number of precursors against Contaminant Repository for Affinity Purification (CRAPome) scores [2]. Identifications with low CRAPome scores and high peptide counts are less likely to be false positive IR interactors. **E**. Scatter plot illustrating the number of precursors identified per protein in the reference proteome against the IR interactome datasets.

**Figure S4.**
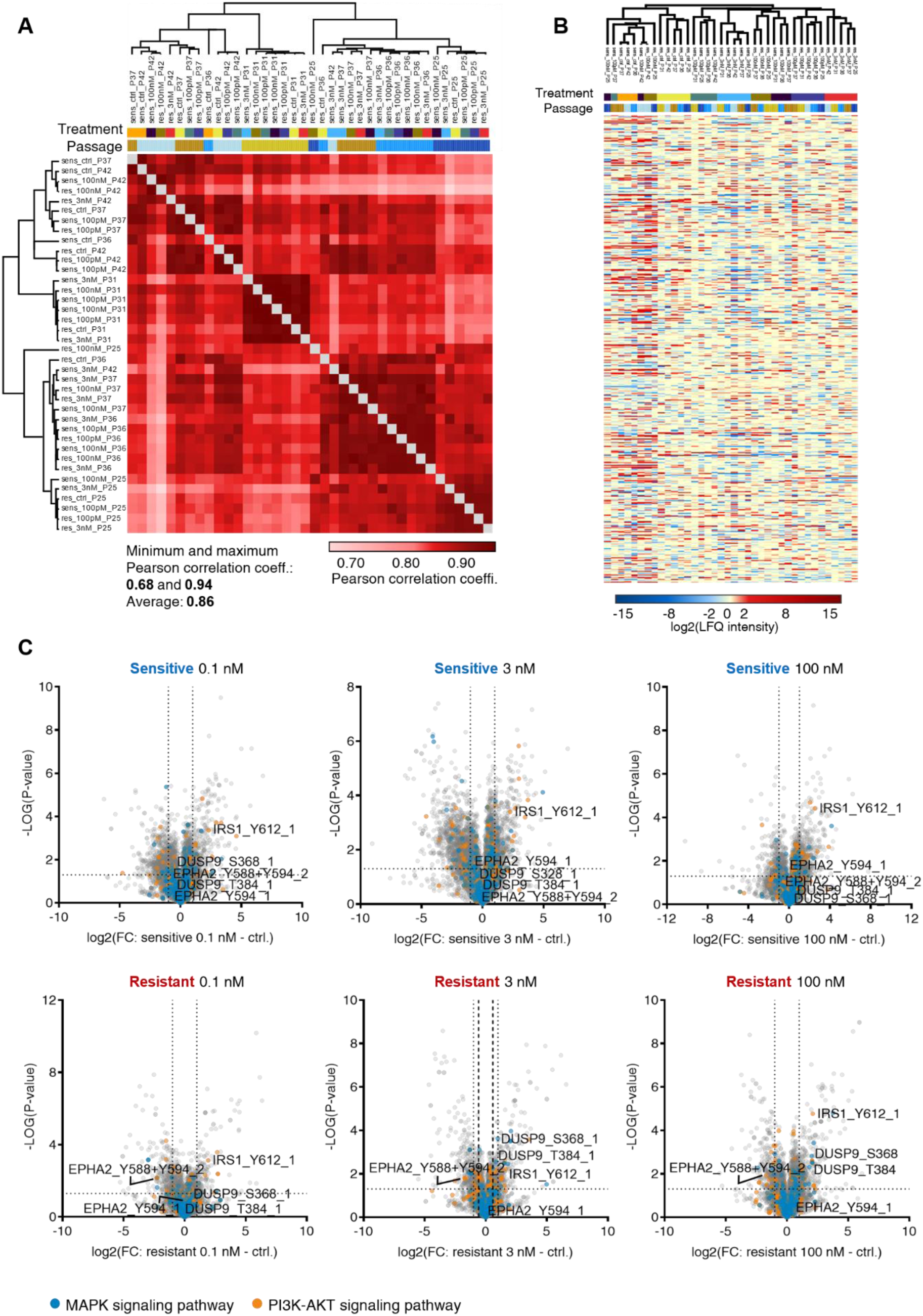
Data quality check of phosphoproteomics covering insulin dose-response in insulin-sensitive and -resistant HepG2 IGF1R KO cells. **A**. Heatmap comparing the Pearson correlation coefficients of the DIA MS-based phosphoproteomics data in insulin-sensitive and insulin-resistant HepG2 IGF1R KO cells. The samples were ordered by hierarchical clustering using the Pearson distance metric algorithm. The heatmap displays the minimum, maximum, and average Pearson correlation coefficients (n=5 biological independent replicates for each condition). **B**. Hierarchical clustering of the log2-transformed LFQ intensities of phosphorylation sites after median subtraction based on cell passages. **C**. Volcano plots showing the -log10(p-value) versus log2(fold change) of phosphorylation site intensities. The fold-change represents insulin stimulation with 0.1, 3, or 100 nM insulin for 5 minutes compared to unstimulated control in insulin-sensitive and -resistant cells (p-value<0.05, 2-fold-change dotted, and 1.5-fold-change dashed line, the latter only shown in 3 nM insulin-resistant). Phosphorylation sites from proteins in MAPK (blue) and PI3K-AKT (orange) signaling pathways are highlighted, and specific DUSP9, EphA2, and IRS1 sites are annotated in all plots.

**Figure S5.**
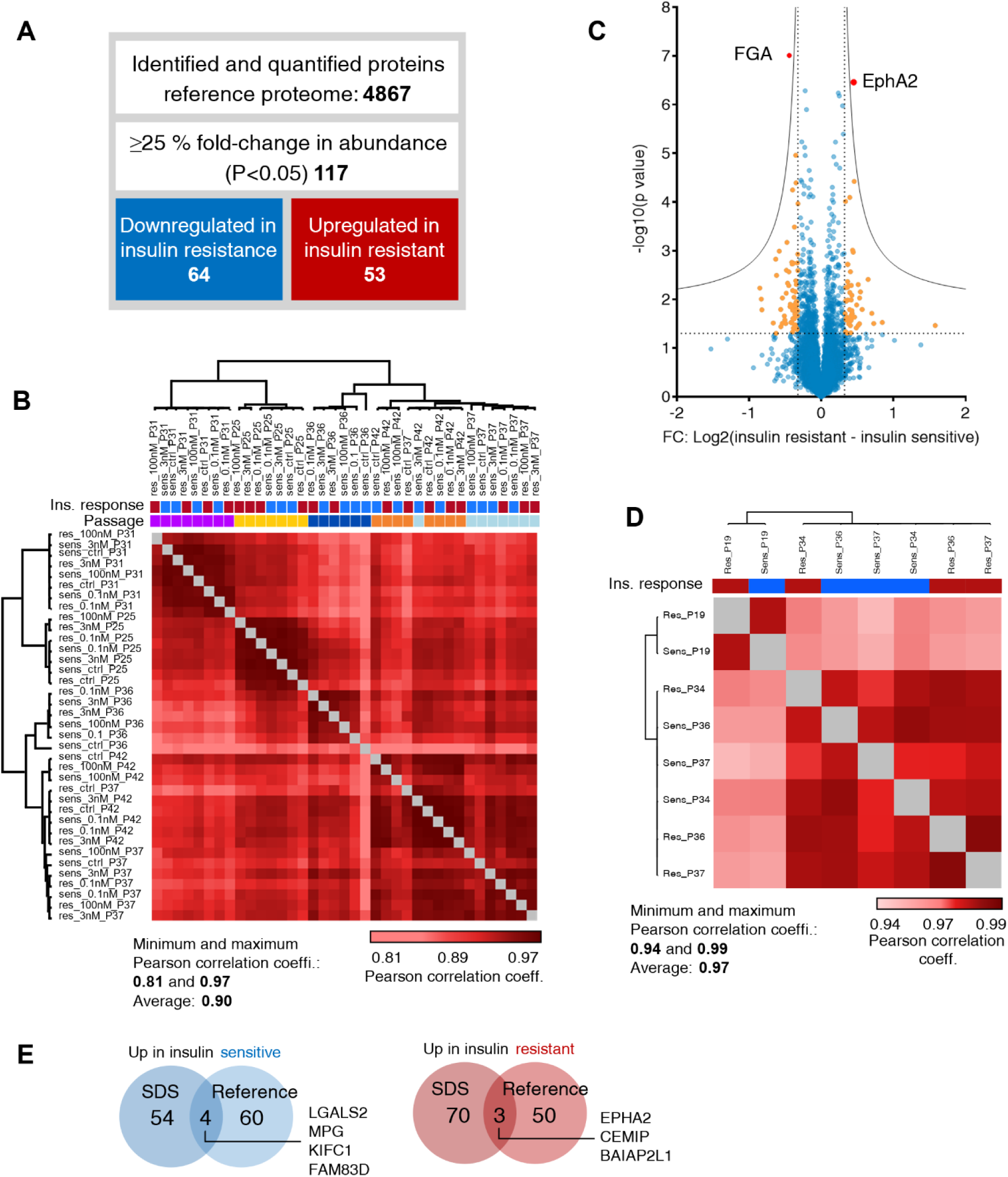
Data quality and comparison of reference and SDS proteome datasets. **A**. Information on the reference proteome data, showing the number of identified proteins and the number of regulated proteins in insulin-sensitive and -resistant cells (≥1.25-fold change and significance, p<0.05). **B**. Heatmap comparing the Pearson correlation coefficients of the DIA MS single-shot proteome in insulin-sensitive and -resistant HepG2 IGF1R KO. The samples were ordered by hierarchical clustering using the Pearson distance metric algorithm with the minimum, maximum, and average Pearson correlation coefficients being stated (n=5 biological independent replicates). **C**. Volcano plot presenting differentially regulated proteins identified in insulin-sensitive and -resistant reference proteome from the single-shot proteome DIA MS analysis. Significantly regulated proteins (≥1.25-fold change and significance, p<0.05 dotted line) are highlighted in orange and (FDR<0.05, S0 = 0.1 solid line) in red with label (n=37). **D**. Same as in B, for the SDS proteome data. **E**. Overlap of proteins identified as being significantly up- or downregulated between insulin-sensitive and -resistant cells in the reference and SDS proteome analyses.

**Figure S6.**
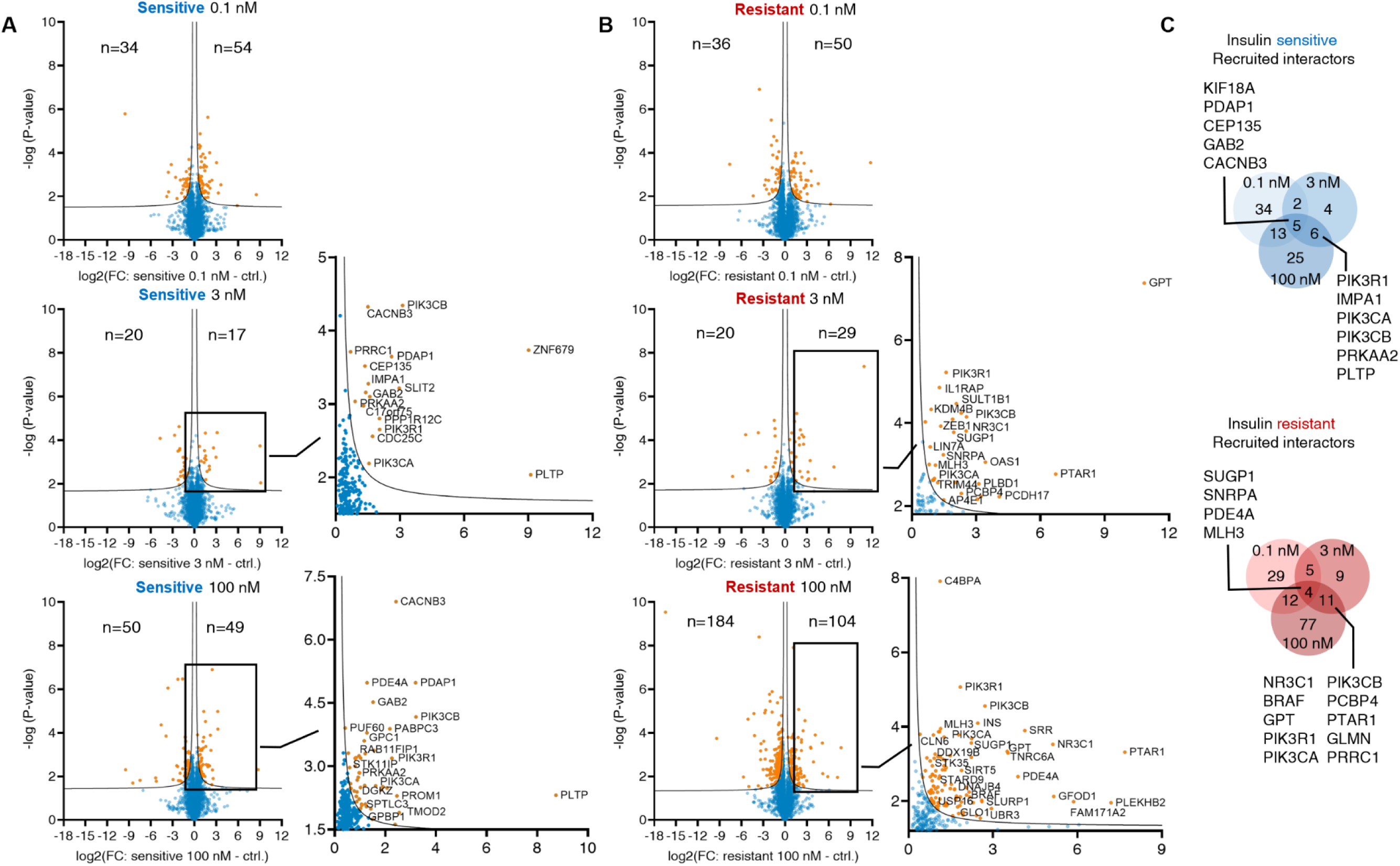
IR interactome response to insulin dose treatment in insulin-sensitive and - resistant HepG2 IGF1R KO cells. Related to Figure 3. **A**. Volcano -log10(p-value) versus log2(fold change) of IR interactor LFQ intensity measured by DIA MS. Illustrating the insulin-stimulated recruited IR interactors, with fold-change difference between unstimulated (ctrl.) and 5-minute 0.1, 3, or 100 nM insulin stimulation, respectively. For the insulin-sensitive cells (two-sided t-test in Perseus software, FDR<0.05, s0=0.1). Zoom on insulin-stimulated interactors upon 5 minutes 3 and 100 nM insulin stimulation. **B**. Same as in A. for the insulin-resistant state. **C**. Overlap of recruited interactors after stimulation with 0.1, 3, and 100 nM insulin. Insulin-sensitive (blue) and -resistant (red) cells.

**Figure S7.**
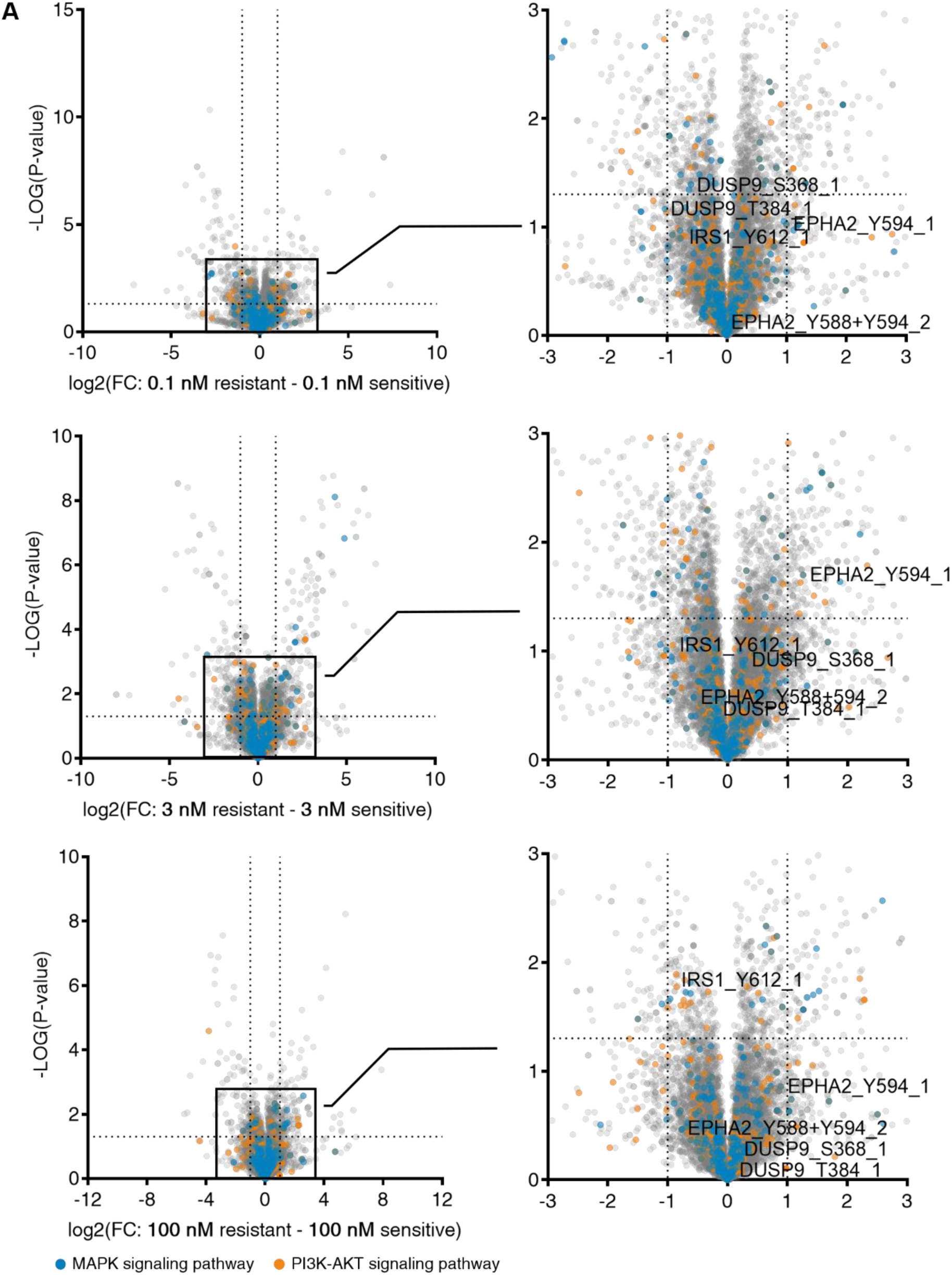
Differential phosphoproteome response in insulin-sensitive and -resistant conditions after stimulation with dosing insulin concentrations. Related to Figure 4. **A**. Volcano plot visualizing -log10(p-value) versus log2(fold change) of phosphorylation site LFQ intensity measured by DIA MS. Fold changes of differentially regulated phospho-sites between insulin-sensitive and -resistant lysates after 5 minutes stimulation with 0.1, 3, or 100 nM insulin. Phosphorylation sites to the right are upregulated in insulin resistance. Phosphorylation sites from MAPK pathway proteins are highlighted in blue and PI3K-AKT pathway proteins in orange. Zoom of each plot to the right with selected phosphorylation sites from DUSP9, EphA2, and IRS1 annotated (dotted line: ≥2-fold change and significance, p<0.05).

**Figure S8.**
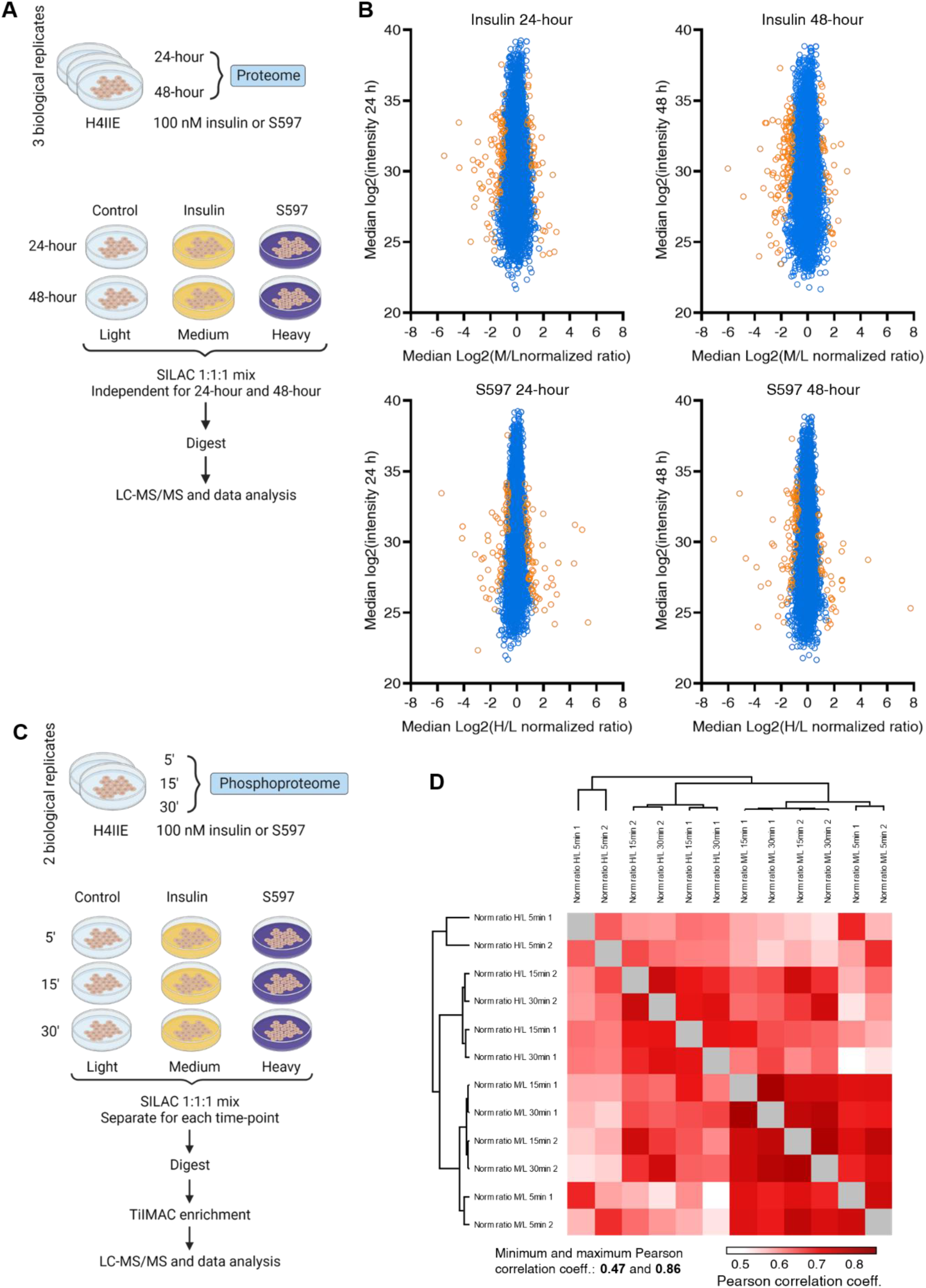
Experimental workflow and data quality control of SILAC MS-data from H4IIE cells. **A**. Experimental design and procedure for SILAC MS-based quantitative proteomics experiment in H4IIE cells treated for 24- or 48-hours with 100 nM insulin or the partial IR agonist S597 (n=3 independent biological replicates). **B**. Plots of log2-transformed summed peptide intensities, as a function of normalized log2-transformed protein ratio for control and insulin or S597 treated cells. Proteins are colored based on p-values for regulation (p<0.05 in orange, and p≥0.05 in blue, from significance B test, with Ben. Ho FDR<0.05 in Perseus). **C**. Experimental design and procedure for generation of SILAC MS-based quantitative phosphoproteomics experiment in H4IIE cells stimulated for 5-, 15-, or 30 minutes with 100 nM insulin or S597 (n=2 independent biological replicates). **D**. Heatmap comparing the Pearson correlation coefficients of the SILAC phosphopeptide ratios in H4IIE after temporal stimulation with 100 nM insulin or S597. The samples were ordered by hierarchical clustering using the Pearson distance metric algorithm. The heatmap displays the minimum, maximum, and average Pearson correlation coefficients.

## References

[1] G. Wilcox, “Insulin and Insulin Resistance,” 2005.

[2] P. N. Freeman AM, *Insulin Resistance*. Treasure Island (FL): StatPearls Publishing, 2023.

[3] V. Parcha et al., “Insulin Resistance and Cardiometabolic Risk Profile among Nondiabetic American Young Adults: Insights from NHANES,” Journal of Clinical Endocrinology and Metabolism, vol. 107, no. 1, pp. E25–E37, Jan. 2022, doi: 10.1210/clinem/dgab645.

[4] A. M. F. Johnson and J. M. Olefsky, “The origins and drivers of insulin resistance,” Cell, vol. 152, no. 4. Elsevier B.V., pp. 673–684, Feb. 14, 2013. doi: 10.1016/j.cell.2013.01.041.

[5] C. J. Nolan and M. Prentki, “Insulin resistance and insulin hypersecretion in the metabolic syndrome and type 2 diabetes: Time for a conceptual framework shift,” Diabetes and Vascular Disease Research, vol. 16, no. 2. SAGE Publications Ltd, pp. 118–127, Mar. 01, 2019. doi: 10.1177/1479164119827611.

[6] J. Boucher, A. Kleinridders, and C. Ronald Kahn, “Insulin receptor signaling in normal and insulin-resistant states,” Cold Spring Harb Perspect Biol, vol. 6, no. 1, Jan. 2014, doi: 10.1101/cshperspect.a009191.

[7] H. C. and M. J. B. Eileen L. Whiteman, “Role of Akt/protein kinase B in metabolism,” Endocrinology & Metabolism, vol. 13, pp. 444–451, Dec. 2002.

[8] A. R. Saltiel, “Insulin signaling in health and disease,” Journal of Clinical Investigation, vol. 131, no. 1. American Society for Clinical Investigation, Jan. 04, 2021. doi: 10.1172/JCI142241.

[9] D. Santoleri and P. M. Titchenell, “Resolving the Paradox of Hepatic Insulin Resistance,” CMGH, vol. 7, no. 2. Elsevier Inc, pp. 447–456, Jan. 01, 2019. doi: 10.1016/j.jcmgh.2018.10.016.

[10] A. Belfiore et al., “Insulin receptor isoforms in physiology and disease: An updated view,” Endocr Rev, vol. 38, no. 5, pp. 379–431, 2017, doi: 10.1210/er.2017-00073.

[11] K. Moelling, K. Schad, M. Bosse, S. Zimmermann, and M. Schweneker, “Regulation of Raf-Akt Cross-talk,” Journal of Biological Chemistry, vol. 277, no. 34, pp. 31099–31106, 2002, doi: 10.1074/jbc.M111974200.

[12] Y. Arkun, “Dynamic Modeling and Analysis of the Cross-Talk between Insulin/AKT and MAPK/ERK Signaling Pathways,” PLoS One, vol. 11, no. 3, pp. e0149684-, Mar. 2016, [Online]. Available: 10.1371/journal.pone.0149684

[13] V. P. Knutson, G. V Ronnett, and M. D. Lane, “Rapid, reversible internalization of cell surface insulin receptors. Correlation with insulin-induced down-regulation.,” Journal of Biological Chemistry, vol. 258, no. 20, pp. 12139–12142, 1983, doi: 10.1016/S0021-9258(17)44146-9.

[14] A. H. Soil, C. R. Kahn, and D. M. Neviile, “Insulin Binding to Liver Plasma Membranes in the Obese Hyperglycemic (o b/o b) Mouse DEMONSTRATION OF A DECREASED NUMBER OF FUNCTIONALLY NORMAL RECEPTORS,” 1975.

[15] J. F. Caro et al., “Studies on the Mechanism of Insulin Resistance in the Liver from Humans with Noninsulin-dependent Diabetes Insulin Action and Binding in Isolated Hepatocytes, Insulin Receptor Structure, and Kinase Activity,” 1986.

[16] L. A. Sechi et al., “Abnormalities of Insulin Receptors in Spontaneously Hypertensive Rats,” Hypertension, vol. 27, no. 4, pp. 955–961, Apr. 1996, doi: 10.1161/01.HYP.27.4.955.

[17] A. Dall’Agnese et al., “The dynamic clustering of insulin receptor underlies its signaling and is disrupted in insulin resistance,” Nat Commun, vol. 13, no. 1, Dec. 2022, doi: 10.1038/s41467-022-35176-7.

[18] K. J. Catalano, B. A. Maddux, J. Szary, J. F. Youngren, I. D. Goldfine, and F. Schaufele, “Insulin resistance induced by hyperinsulinemia coincides with a persistent alteration at the insulin receptor tyrosine kinase domain,” PLoS One, vol. 9, no. 9, Sep. 2014, doi: 10.1371/journal.pone.0108693.

[19] C. Francavilla et al., “Multilayered proteomics reveals molecular switches dictating ligand-dependent EGFR trafficking,” Nat Struct Mol Biol, vol. 23, no. 6, pp. 608–618, Jun. 2016, doi: 10.1038/nsmb.3218.

[20] K. B. Emdal et al., “Temporal proteomics of NGF-TrkA signaling identifies an inhibitory role for the E3 ligase Cbl-b in neuroblastoma cell differentiation,” Sci Signal, vol. 8, no. 374, pp. ra40–ra40, Apr. 2015, doi: 10.1126/scisignal.2005769.

[21] D. J. Fazakerley et al., “Phosphoproteomics reveals rewiring of the insulin signaling network and multi-nodal defects in insulin resistance,” Nat Commun, vol. 14, no. 1, p. 923, 2023, doi: 10.1038/s41467-023-36549-2.

[22] R. Lennon, S. B. Hosawi, J. D. Humphries, R. J. Coward, D. Knight, and M. J. Humphries, “Global proteomic analysis of insulin receptor interactors in glomerular podocytes,” Wellcome Open Res, vol. 5, 2020, doi: 10.12688/wellcomeopenres.16072.1.

[23] K. Salokas et al., “Physical and functional interactome atlas of human receptor tyrosine kinases,” EMBO Rep, vol. 23, no. 6, Jun. 2022, doi: 10.15252/embr.202154041.

[24] E. L. Huttlin et al., “Dual proteome-scale networks reveal cell-specific remodeling of the human interactome,” Cell, vol. 184, no. 11, pp. 3022–3040.e28, 2021, doi: 10.1016/j.cell.2021.04.011.

[25] H. Miao et al., “EphA2 Mediates Ligand-Dependent Inhibition and Ligand-Independent Promotion of Cell Migration and Invasion via a Reciprocal Regulatory Loop with Akt,” Cancer Cell, vol. 16, no. 1, pp. 9–20, Jul. 2009, doi: 10.1016/j.ccr.2009.04.009.

[26] J. E. Park, A. I. Son, and R. Zhou, “Roles of EphA2 in development and disease,” Genes, vol. 4, no. 3. pp. 334–357, Sep. 2013. doi: 10.3390/genes4030334.

[27] T. Xiao, Y. Xiao, W. Wang, Y. Y. Tang, Z. Xiao, and M. Su, “Targeting EphA2 in cancer,” Journal of Hematology and Oncology, vol. 13, no. 1. BioMed Central, Aug. 18, 2020. doi: 10.1186/s13045-020-00944-9.

[28] M. Tandon, S. V. Vemula, and S. K. Mittal, “Emerging strategies for EphA2 receptor targeting for cancer therapeutics,” Expert Opinion on Therapeutic Targets, vol. 15, no. 1. pp. 31–51, Jan. 2011. doi: 10.1517/14728222.2011.538682.

[29] M. Jensen, B. Hansen, P. De Meyts, L. Schäffer, and B. Ursø, “Activation of the Insulin Receptor by Insulin and a Synthetic Peptide Leads to Divergent Metabolic and Mitogenic Signaling and Responses,” Journal of Biological Chemistry, vol. 282, no. 48, pp. 35179– 35186, 2007, doi: 10.1074/jbc.M704599200.

[30] M. Jensen, J. Palsgaard, R. Borup, P. De Meyts, and L. Schäffer, “Activation of the insulin receptor (IR) by insulin and a synthetic peptide has different effects on gene expression in IR-transfected L6 myoblasts,” Biochemical Journal, vol. 412, no. 3, pp. 435–445, Jun. 2008, doi: 10.1042/BJ20080279.

[31] K. Rufinatscha et al., “Metabolic effects of reduced growth hormone action in fatty liver disease,” Hepatol Int, vol. 12, no. 5, pp. 474–481, Sep. 2018, doi: 10.1007/s12072-018-9893-7.

[32] R. D. Kineman, M. del Rio-Moreno, and A. Sarmento-Cabral, “40 years of IGF1: Understanding the tissue-specific roles of IGF1/IGF1R in regulating metabolism using the Cre/loxP system,” Journal of Molecular Endocrinology, vol. 61, no. 1. BioScientifica Ltd., pp. T187–T198, Jul. 01, 2018. doi: 10.1530/JME-18-0076.

[33] M. A. Soos et al., “Monoclonal antibodies reacting with multiple epitopes on the human insulin receptor,” 1986.

[34] L. H. Ørstrup et al., “Cross-species reactive monoclonal antibodies against the extracellular domains of the insulin receptor and IGF1 receptor,” J Immunol Methods, vol. 465, pp. 20–26, 2019, doi: 10.1016/j.jim.2018.11.014.

[35] S. A. Bustin et al., “The MIQE Guidelines: Minimum Information for Publication of Quantitative Real-Time PCR Experiments,” Clin Chem, vol. 55, no. 4, pp. 611–622, Apr. 2009, doi: 10.1373/clinchem.2008.112797.

[36] T. S. Batth et al., “Protein aggregation capture on microparticles enables multipurpose proteomics sample preparation,” Molecular and Cellular Proteomics, vol. 18, no. 5, pp. 1027–1035, May 2019, doi: 10.1074/mcp.TIR118.001270.

[37] D. B. Bekker-Jensen et al., “Rapid and site-specific deep phosphoproteome profiling by data-independent acquisition without the need for spectral libraries,” Nat Commun, vol. 11, no. 1, Dec. 2020, doi: 10.1038/s41467-020-14609-1.

[38] C. Koenig, A. Martinez-Val, P. Naicker, S. Stoychev, J. Jordaan, and J. V Olsen, “Protocol for high-throughput semi-automated label-free- or TMT-based phosphoproteome profiling,” STAR Protoc, vol. 4, no. 3, p. 102536, 2023, doi: 10.1016/j.xpro.2023.102536.

[39] T. S. Batth, C. Francavilla, and J. V Olsen, “Off-Line High-pH Reversed-Phase Fractionation for In-Depth Phosphoproteomics,” J Proteome Res, vol. 13, no. 12, pp. 6176–6186, Dec. 2014, doi: 10.1021/pr500893m.

[40] M. R. Larsen, T. E. Thingholm, O. N. Jensen, P. Roepstorff, and T. J. D. Jørgensen, “Highly Selective Enrichment of Phosphorylated Peptides from Peptide Mixtures Using Titanium Dioxide Microcolumns*,” Molecular & Cellular Proteomics, vol. 4, no. 7, pp. 873– 886, 2005, doi: 10.1074/mcp.T500007-MCP200.

[41] M. W. H. Pinkse, P. M. Uitto, M. J. Hilhorst, B. Ooms, and A. J. R. Heck, “Selective Isolation at the Femtomole Level of Phosphopeptides from Proteolytic Digests Using 2D-NanoLC-ESI-MS/MS and Titanium Oxide Precolumns,” Anal Chem, vol. 76, no. 14, pp. 3935–3943, Jul. 2004, doi: 10.1021/ac0498617.

[42] R. Bruderer et al., “Extending the Limits of Quantitative Proteome Profiling with Data-Independent Acquisition and Application to Acetaminophen-Treated Three-Dimensional Liver Microtissues*[S],” Molecular & Cellular Proteomics, vol. 14, no. 5, pp. 1400–1410, 2015, doi: 10.1074/mcp.M114.044305.

[43] J. Cox and M. Mann, “MaxQuant enables high peptide identification rates, individualized p.p.b.-range mass accuracies and proteome-wide protein quantification,” Nat Biotechnol, vol. 26, no. 12, pp. 1367–1372, 2008, doi: 10.1038/nbt.1511.

[44] S. Wieczorek et al., “DAPAR & ProStaR: software to perform statistical analyses in quantitative discovery proteomics,” Bioinformatics, vol. 33, no. 1, pp. 135–136, Jan. 2017, doi: 10.1093/bioinformatics/btw580.

[45] D. Mellacheruvu et al., “The CRAPome: A contaminant repository for affinity purification-mass spectrometry data,” Nat Methods, vol. 10, no. 8, pp. 730–736, Aug. 2013, doi: 10.1038/nmeth.2557.

[46] J. J. Almagro Armenteros, C. K. Sønderby, S. K. Sønderby, H. Nielsen, and O. Winther, “DeepLoc: prediction of protein subcellular localization using deep learning,” Bioinformatics, vol. 33, no. 21, pp. 3387–3395, Nov. 2017, doi: 10.1093/bioinformatics/btx431.

[47] S. Tyanova et al., “The Perseus computational platform for comprehensive analysis of (prote)omics data,” Nature Methods, vol. 13, no. 9. Nature Publishing Group, pp. 731–740, Aug. 30, 2016. doi: 10.1038/nmeth.3901.

[48] D. Szklarczyk et al., “The STRING database in 2023: protein–protein association networks and functional enrichment analyses for any sequenced genome of interest,” Nucleic Acids Res, vol. 51, no. D1, pp. D638–D646, Jan. 2023, doi: 10.1093/nar/gkac1000.

[49] P. Shannon et al., “Cytoscape: A software Environment for integrated models of biomolecular interaction networks,” Genome Res, vol. 13, no. 11, pp. 2498–2504, Nov. 2003, doi: 10.1101/gr.1239303.

[50] K. Breuer et al., “InnateDB: Systems biology of innate immunity and beyond - Recent updates and continuing curation,” Nucleic Acids Res, vol. 41, no. D1, Jan. 2013, doi: 10.1093/nar/gks1147.

[51] J. V. Olsen et al., “Global, In Vivo, and Site-Specific Phosphorylation Dynamics in Signaling Networks,” Cell, vol. 127, no. 3, pp. 635–648, Nov. 2006, doi: 10.1016/j.cell.2006.09.026.

[52] Y. Perez-Riverol et al., “The PRIDE database and related tools and resources in 2019: improving support for quantification data,” Nucleic Acids Res, vol. 47, no. D1, pp. D442– D450, Jan. 2019, doi: 10.1093/nar/gky1106.

[53] R. D. Kineman, M. del Rio-Moreno, and A. Sarmento-Cabral, “40 years of IGF1: Understanding the tissue-specific roles of IGF1/IGF1R in regulating metabolism using the Cre/loxP system,” Journal of Molecular Endocrinology, vol. 61, no. 1. BioScientifica Ltd., pp. T187–T198, Jul. 01, 2018. doi: 10.1530/JME-18-0076.

[54] J. R. Gavin, J. Roth, D. M. Neville, P. De Meyts, and D. N. Buellt, “Insulin-Dependent Regulation of Insulin Receptor Concentrations: A Direct Demonstration in Cell Culture,” 1974. [Online]. Available: https://www.pnas.org

[55] C. Hall, H. Yu, and E. Choi, “Insulin receptor endocytosis in the pathophysiology of insulin resistance,” Experimental and Molecular Medicine, vol. 52, no. 6. Springer Nature, pp. 911–920, Jun. 01, 2020. doi: 10.1038/s12276-020-0456-3.

[56] X. Liu et al., “Insulin induces insulin receptor degradation in the liver through EphB4,” Nat Metab, vol. 4, no. 9, pp. 1202–1213, Sep. 2022, doi: 10.1038/s42255-022-00634-5.

[57] I. Wittig and H. Schägger, “Native electrophoretic techniques to identify protein-protein interactions,” Proteomics, vol. 9, no. 23. Wiley-VCH Verlag, pp. 5214–5223, 2009. doi: 10.1002/pmic.200900151.

[58] N. Nagaraj et al., “Deep proteome and transcriptome mapping of a human cancer cell line,” Mol Syst Biol, vol. 7, 2011, doi: 10.1038/msb.2011.81.

[59] J. C. Schafer, R. E. McRae, E. H. Manning, L. A. Lapierre, and J. R. Goldenring, “Rab11-FIP1A regulates early trafficking into the recycling endosomes,” Exp Cell Res, vol. 340, no. 2, pp. 259–273, 2016, doi: 10.1016/j.yexcr.2016.01.003.

[60] Z. Qu, S. Ji, and S. Zheng, “Braf controls the effects of metformin on neuroblast cell divisions in c. Elegans,” Int J Mol Sci, vol. 22, no. 1, pp. 1–13, Jan. 2021, doi: 10.3390/ijms22010178.

[61] O. J. Kennedy et al., “Prognostic and predictive value of metformin in the European Organisation for Research and Treatment of Cancer 1325/KEYNOTE-054 phase III trial of pembrolizumab versus placebo in resected high-risk stage III melanoma,” Eur J Cancer, vol. 189, p. 112900, 2023, doi: 10.1016/j.ejca.2023.04.016.

[62] R. R. Henry, B. Gumbiner, T. Flynn, and A. W. Thorburn, “Metabolic Effects of Hyperglycemia and Hyperinsulinemia on Fate of Intracellular Glucose in NIDDM,” 1990. [Online]. Available: http://diabetesjournals.org/diabetes/article-pdf/39/2/149/357273/39-2-149.pdf

[63] R. A. Vaidya et al., “Hyperinsulinemia: an early biomarker of metabolic dysfunction,” Frontiers in Clinical Diabetes and Healthcare, vol. 4, May 2023, doi: 10.3389/fcdhc.2023.1159664.

[64] R. Amanchy, B. Periaswamy, S. Mathivanan, R. Reddy, S. G. Tattikota, and A. Pandey, “A curated compendium of phosphorylation motifs,” Nat Biotechnol, vol. 25, no. 3, pp. 285–286, 2007, doi: 10.1038/nbt0307-285.

[65] F. Z. Khoubai and C. F. Grosset, “Dusp9, a dual-specificity phosphatase with a key role in cell biology and human diseases,” International Journal of Molecular Sciences, vol. 22, no. 21. MDPI, Nov. 01, 2021. doi: 10.3390/ijms222111538.

[66] B. Emanuelli, D. Eberlé, R. Suzuki, and C. R. Kahn, “Overexpression of the dual-specificity phosphatase MKP-4/DUSP-9 protects against stress-induced insulin resistance,” Proceedings of the National Academy of Sciences, vol. 105, no. 9, pp. 3545– 3550, Mar. 2008, doi: 10.1073/pnas.0712275105.

[67] S. Y. Yoon, J. H. Lee, S. J. Kwon, H. J. Kang, and S. J. Chung, “Ginkgolic acid as a dual-targeting inhibitor for protein tyrosine phosphatases relevant to insulin resistance,” Bioorg Chem, vol. 81, pp. 264–269, Dec. 2018, doi: 10.1016/j.bioorg.2018.08.011.

[68] W. J. A. J. Hendriks and R. Pulido, “Protein tyrosine phosphatase variants in human hereditary disorders and disease susceptibilities,” Biochimica et Biophysica Acta - Molecular Basis of Disease, vol. 1832, no. 10. pp. 1673–1696, Oct. 2013. doi: 10.1016/j.bbadis.2013.05.022.

[69] K. P. Hoeflich et al., “Intermittent Administration of MEK Inhibitor GDC-0973 plus PI3K Inhibitor GDC-0941 Triggers Robust Apoptosis and Tumor Growth Inhibition,” Cancer Res, vol. 72, no. 1, pp. 210–219, Jan. 2012, doi: 10.1158/0008-5472.CAN-11-1515.

[70] E. Agarwal, A. Chaudhuri, P. D. Leiphrakpam, K. L. Haferbier, M. G. Brattain, and S. Chowdhury, “Akt inhibitor MK-2206 promotes anti-tumor activity and cell death by modulation of AIF and Ezrin in colorectal cancer,” BMC Cancer, vol. 14, no. 1, p. 145, 2014, doi: 10.1186/1471-2407-14-145.

[71] L. Yan, “Abstract #DDT01-1: MK-2206: A potent oral allosteric AKT inhibitor,” Cancer Res, vol. 69, no. 9_Supplement, pp. DDT01-1-DDT01-1, May 2009.

[72] H. Hvid et al., “Increased insulin receptor binding and increased IGF-1 receptor binding are linked with increased growth of L6hIR cell xenografts in vivo,” Sci Rep, vol. 10, no. 1, Dec. 2020, doi: 10.1038/s41598-020-64318-4.

[73] W. Cai et al., “Domain-dependent effects of insulin and IGF-1 receptors on signalling and gene expression,” Nat Commun, vol. 8, no. 1, p. 14892, 2017, doi: 10.1038/ncomms14892.

[74] R. Lammers, A. Gray, J. Schlessinger, and A. Ullrich, “Differential signalling potential of insulin- and IGF-1-receptor cytoplasmic domains,” EMBO Journal, vol. 8, no. 5, pp. 1369– 1375, 1989, doi: 10.1002/j.1460-2075.1989.tb03517.x.

[75] R. Schumacher, L. Mosthaf, J. Schlessinger, D. Brandenburg, and A. Ullrich, “Insulin and insulin-like growth factor-1 binding specificity is determined by distinct regions of their cognate receptors,” Journal of Biological Chemistry, vol. 266, no. 29, pp. 19288–19295, 1991, doi: 10.1016/s0021-9258(18)54996-6.

[76] R. Slaaby et al., “Hybrid Receptors Formed by Insulin Receptor (IR) and Insulin-like Growth Factor I Receptor (IGF-IR) Have Low Insulin and High IGF-1 Affinity Irrespective of the IR Splice Variant*,” Journal of Biological Chemistry, vol. 281, no. 36, pp. 25869– 25874, 2006, doi: 10.1074/jbc.M605189200.

[77] M. Federici et al., “Increased expression of insulin/insulin-like growth factor-1 hybrid receptors in skeletal muscle of noninsulin-dependent diabetes mellitus subjects,” Journal of Clinical Investigation, vol. 98, no. 12, pp. 2887–2893, Dec. 1996, doi: 10.1172/JCI119117.

[78] K. Cusi et al., “Insulin resistance differentially affects the PI 3-kinase– and MAP kinase– mediated signaling in human muscle,” J Clin Invest, vol. 105, no. 3, 1999.

[79] B. Draznin, “Mechanism of the mitogenic influence of hyperinsulinemia,” Diabetology and Metabolic Syndrome, vol. 3, no. 1. 2011. doi: 10.1186/1758-5996-3-10.

[80] S. Emamgholipour, R. Ebrahimi, A. Bahiraee, F. Niazpour, and R. Meshkani, “Acetylation and insulin resistance: a focus on metabolic and mitogenic cascades of insulin signaling,” Crit Rev Clin Lab Sci, vol. 57, no. 3, pp. 196–214, Apr. 2020, doi: 10.1080/10408363.2019.1699498.

[81] K. Ozaki et al., “Targeting the ERK signaling pathway as a potential treatment for insulin resistance and type 2 diabetes,” American Journal of Physiology-Endocrinology and Metabolism, vol. 310, no. 8, pp. E643–E651, Feb. 2016, doi: 10.1152/ajpendo.00445.2015.

[82] P. Ye et al., “Dual-specificity Phosphatase 9 Protects Against Non-alcoholic Fatty Liver Disease in Mice via ASK1 Suppression,” Hepatology, vol. 69, Jul. 2018, doi: 10.1002/hep.30198.

[83] H. F. Chen, H. C. Chuang, and T. H. Tan, “Regulation of dual-specificity phosphatase (Dusp) ubiquitination and protein stability,” International Journal of Molecular Sciences, vol. 20, no. 11. MDPI AG, Jun. 01, 2019. doi: 10.3390/ijms20112668.

[84] P. Dong, J.-Y. Zhou, and G. S. Wu, “Post-translational regulation of mitogen-activated protein kinase phosphatase-2 (MKP-2) by ERK,” Cell Cycle, vol. 9, no. 23, pp. 4650– 4655, Dec. 2010, doi: 10.4161/cc.9.23.13957.

[85] M. Macrae et al., “A conditional feedback loop regulates Ras activity through EphA2,” Cancer Cell, vol. 8, no. 2, pp. 111–118, 2005, doi: 10.1016/j.ccr.2005.07.005.

[86] M. Federici et al., “Increased Abundance of Insulin/Insulin-Like Growth Factor-I Hybrid Receptors in Skeletal Muscle of Obese Subjects Is Correlated with In Vivo Insulin Sensitivity*,” 1998. [Online]. Available: https://academic.oup.com/jcem/article/83/8/2911/2660616

[87] M. A. Soos, C. E. Field, and K. Siddle, “Purified hybrid insulin/insulin-like growth factor-I receptors bind insulin-like growth factor-I, but not insulin, with high affinity,” 1993.

